# PrkA controls peptidoglycan biosynthesis through the essential phosphorylation of ReoM

**DOI:** 10.1101/2019.12.16.877605

**Authors:** Sabrina Wamp, Zoe J. Rutter, Jeanine Rismondo, Claire E. Jennings, Lars Möller, Richard J. Lewis, Sven Halbedel

## Abstract

Peptidoglycan (PG) is the main component of bacterial cell walls and the target for many antibiotics. PG biosynthesis is tightly coordinated with cell wall growth and turnover, and many of these control activities depend upon PASTA-domain containing eukaryotic-like serine/threonine protein kinases (PASTA-eSTK) that sense PG fragments. However, only a few PG biosynthetic enzymes are direct kinase substrates. Here, we identify the conserved ReoM protein as a novel PASTA-eSTK substrate in the Gram-positive pathogen *Listeria monocytogenes*. Our data show that the phosphorylation of ReoM is essential as it controls ClpCP-dependent proteolytic degradation of the essential enzyme MurA, which catalyses the first committed step in PG biosynthesis. We also identify ReoY as a second novel factor required for degradation of ClpCP substrates. Collectively, our data imply that the first committed step of PG biosynthesis is activated through control of ClpCP protease activity in response to signals of PG homeostasis imbalance.

## INTRODUCTION

The cell wall of Gram-positive bacteria is a complicated three-dimensional structure that engulfs the cell as a closed sacculus. The main component of bacterial cell walls is peptidoglycan (PG), a network of glycan strands crosslinked together by short peptides (1). PG biosynthesis starts with the conversion of UDP-Glc*N*Ac into lipid II, a disaccharide pentapeptide that is ligated to a membrane-embedded bactoprenol carrier lipid (2). This monomeric PG precursor is then flipped from the inner to the outer leaflet of the cytoplasmic membrane by MurJ- and Amj-like enzymes called flippases (3–5). Glycosyltransferases belonging either to the bifunctional <u>p</u>enicillin <u>b</u>inding <u>p</u>roteins (PBPs) or the SEDS (<u>s</u>hape, <u>e</u>longation, <u>d</u>ivision and <u>s</u>porulation) family, then transfer the disaccharide pentapeptides to growing PG strands, which are finally crosslinked by a transpeptidation reaction catalysed by monofunctional (class B) or bifunctional (class A) PBPs (6–9). Numerous hydrolytic or PG-modifying enzymes are also required to adapt the sacculus to the morphological changes that occur during bacterial cell growth and division (10, 11) or to alter its chemical properties for instance for immune evasion (12). A suite of regulators ensure that spatiotemporal control of PG synthesis is balanced against PG hydrolysis in cycles of bacterial growth and division (13).

The activity of several key enzymes along the PG biosynthetic pathway is regulated by PASTA (<u>P</u>BP <u>a</u>nd <u>s</u>erine/threonine kinase <u>a</u>ssociated) domain-containing <u>e</u>ukaryotic-like <u>s</u>erine/threonine protein <u>k</u>inases (PASTA-eSTKs) (14–16). These membrane-integral enzymes comprise a cytoplasmic kinase domain linked to several extracellular PASTA domains (15). These proteins are stimulated by free muropeptides and lipid II (that accumulate during damage and turnover of PG) on interaction with their PASTA domains (17–19). PknB, a representative PASTA-eSTK from *Mycobacterium tuberculosis*, phosphorylates GlmU, a bifunctional uridyltransferase/acetyltransferase important for synthesis of UDP-Glc*N*Ac, and in so doing reduces GlmU activity (20). *M. tuberculosis* MviN, a MurJ-like flippase, is also a substrate of PknB and, in its phosphorylated state, P-MviN is inhibited by its binding partner, FhaA (21). *M. tuberculosis* PknB also phosphorylates both the class A PBP PonA1 (22) and the amidase-like PG-hydrolase CwlM, which is essential for growth (23–25). CwlM is membrane-associated and interacts with MurJ to control lipid II export (25). However, when phosphorylated, P-CwlM re-locates from the membrane to the cytoplasm (25) where it allosterically activates MurA 20–40-fold (24). MurA catalyzes the first committed step of PG biosynthesis by transferring an enoylpyruvate moiety to UDP-Glc*N*Ac; MurA is essential in *M. tuberculosis* and in many other bacterial species tested (26–29). Finally, the *Listeria monocytogenes* PASTA-eSTK, PrkA, phosphorylates YvcK, which is required for cell wall homeostasis in a so far unknown way (30). Numerous additional proteins acting to coordinate cell wall biosynthesis with cell division are substrates of PASTA-eSTKs in other Gram-positive bacteria (15), including the late cell division protein GpsB of *Bacillus subtilis* (31, 32). We have shown previously that GpsB from *L. monocytogenes* is important for the last two steps of PG biosynthesis, *i. e.* transglycosylation and transpeptidation, by providing an assembly platform for the class A PBP, PBP A1 (33–36), and this adaptor function of GpsB is maintained in at least *B. subtilis* and *Streptococcus pneumoniae* (35). An *L. monocytogenes* Δ*gpsB* mutant is impaired in PG biosynthesis and cannot grow at elevated temperatures (33), but this phenotype is readily corrected by a suppressor mutation, which mapped to *clpC* (29). ClpC is the ATPase subunit of the ClpCP protease that degrades substrate proteins upon heat stress (37). MurA (*aka* MurAA in *B. subtilis*) is a ClpCP substrate in both *B. subtilis* and *L. monocytogenes* (27, 29) and strongly accumulates in a *L. monocytogenes* Δ*clpC* mutant (29). Thus, a deficiency in the final two enzymatic steps of PG biosynthesis in the absence of GpsB is corrected by mutations in *clpC* that increase the amount of the first enzyme of the same PG biosynthetic pathway.

We here have isolated further *gpsB* suppressor mutations affecting previously unstudied *Listeria* genes. We demonstrate that these proteins control the ClpCP-dependent degradation of MurA in a PrkA-dependent and hitherto unprecedented manner. One of them is phosphorylated by PrkA and this phosphorylation is essential. Our results represent the first molecular link between PrkA-dependent protein phosphorylation and control of PG production in low G/C Gram-positive bacteria and explain how PG biosynthesis is adjusted in these bacteria to meet PG production and repair needs.

## RESULTS

### *gpsB* suppressor mutations in the *lmo1503* (*reoM*) and *lmo1921* (*reoY*) genes

A *L. monocytogenes* Δ*gpsB* mutant is unable to replicate at 42°C, but readily forms suppressors correcting this defect (29). Previously isolated *gpsB* suppressors carried a mutation in the *clpC* gene, important for the stability of the UDP-*N*-acetylglucosamine 1-carboxyvinyltransferase MurA (29). We have characterized three more *shg* (<u>s</u>uppression of <u>h</u>eat sensitive <u>g</u>rowth) suppressor mutants (*shg8*, *shg10* and *shg12*) isolated from a Δ*gpsB* mutant incubated on a BHI agar plate at 42°C. These three *shg* strains grew as fast as the wild type when cultivated at 37°C or 42°C, whereas the parental Δ*gpsB* mutant grew at a reduced rate at 37°C and did not grow at 42°C (Fig. 1A-B), as shown previously (33).

**Figure 1:**
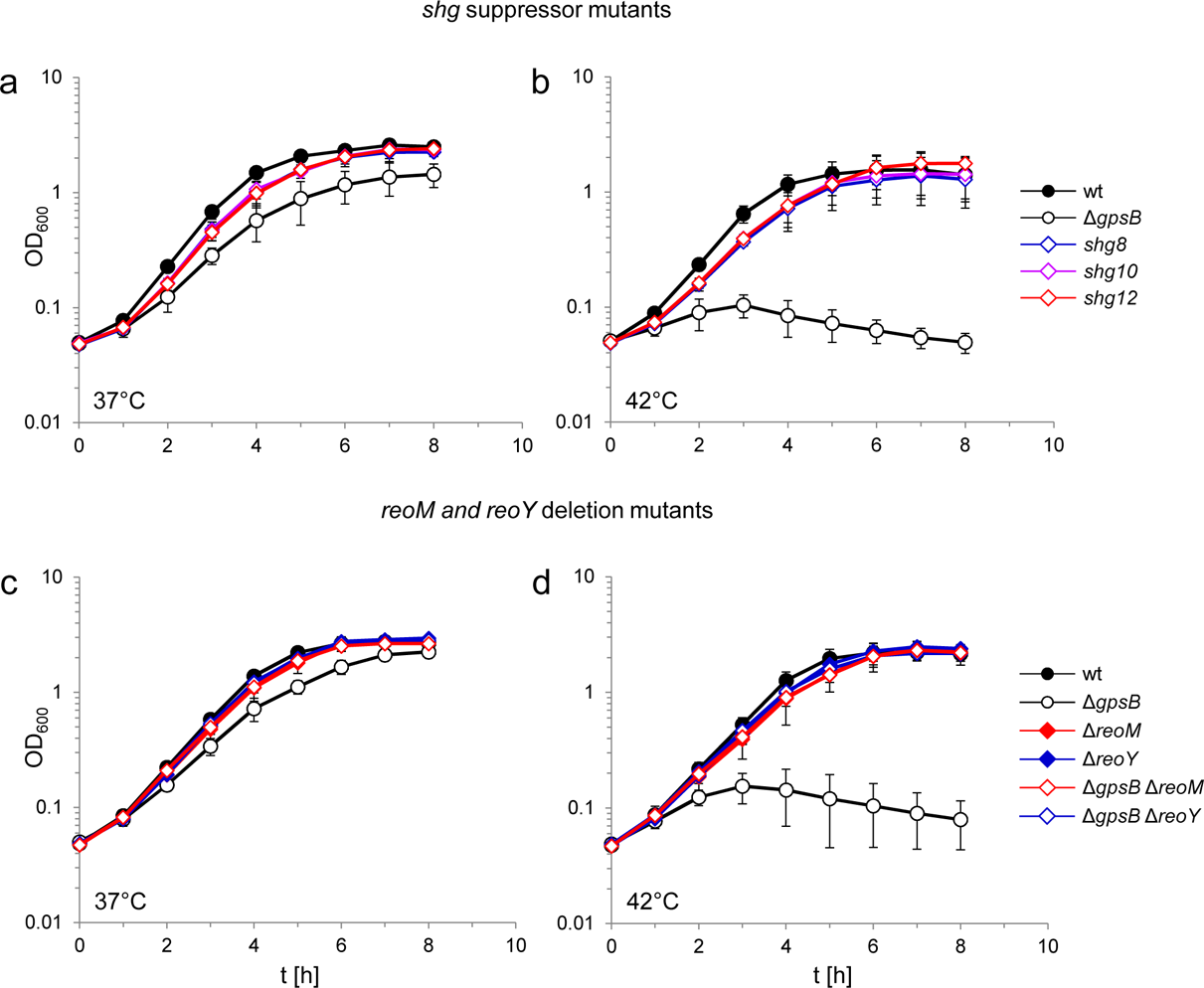
Suppression of the growth defects of a *L. monocytogenes* Δ*gpsB* mutant by *reoM* and *reoY* mutations. (A-B) Effect of suppressor mutations on growth of the Δ*gpsB* mutant. Growth of *L. monocytogenes* strains EGD-e (wt), LMJR19 (Δ*gpsB*), *shg8* (Δ*gpsB reoY* H87Y), *shg10* (Δ*gpsB reoY* TAA74) and *shg12* (Δ*gpsB reoM* RBS mutation) in BHI broth at 37°C (A) and 42°C (B). (C-D) Effect of Δ*reoM* and Δ*reoY* deletions on growth of *L. monocytogenes.* Growth of *L. monocytogenes* strains EGD-e (wt), LMJR19 (Δ*gpsB*), LMSW30 (Δ*reoM*), LMSW32 (Δ*reoY*), LMJR137 (Δ*gpsB* Δ*reoM*) and LMJR120 (Δ*gpsB* Δ*reoY*) in BHI broth was recorded at 37°C (C) and 42°C (D). All growth experiments were performed three times and average values and standard deviations are shown.

Sequencing of the *shg8*, *shg10* and *shg12* genomes identified one SNP in each strain that was absent from the parental Δ*gpsB* mutant. Strain *shg8* carried a mutation in the uncharacterized *lmo1921* gene (herein named *reoY*, see below) that exchanged H87 into tyrosine; the same gene was affected by the introduction of a premature stop codon after the 73^rd^ *reoY* codon in strain *shg10*. Strain *shg12* carried a mutation in the ribosomal binding site (RBS) of the *lmo1503* gene (renamed *reoM*), encoding an IreB-like protein, the function of which is not understood (38). Whether the mutation in the RBS of *reoM* in strain *shg12* affected *reoM* expression, was not clear. Therefore, the *reoM* gene was deleted from the genome of the wild type and the Δ*gpsB* mutant. While deletion of *reoM* had no effect on growth of wild type bacteria, it completely suppressed the growth defects of the Δ*gpsB* mutant at both 37°C and 42°C (Fig. 1C-D). It is thus likely that the mutation in the *reoM* RBS impairs its expression. Likewise, deletion of *reoY* completely restored growth of the Δ*gpsB* mutant at both temperatures (Fig. 1C-D).

Expression of an additional, plasmid-borne copy of *reoM* impaired growth of the Δ*gpsB* mutant without affecting the growth of wild type bacteria, whilst expression of a second copy of *reoY* had no effect (Fig. S1A,B). The expression of *reoM* is thus inversely correlated with the growth of the Δ*gpsB* mutant. Finally, the physiology of the Δ*reoM* and Δ*reoY* mutants was examined; their cell lengths were wild type-like and unaffected by the presence or absence of *gpsB*, suggesting the absence of cell division defects in the Δ*reoM* or Δ*reoY* mutants (Fig. S2A,B). Scanning electron micrographs of Δ*reoM* and Δ*reoY* single mutants revealed that these bacteria had a normal rod-shape, but that the Δ*gpsB* Δ*reoM* and Δ*gpsB* Δ*reoY* double mutants were partially bent (Fig. S2C), implying the presence of some shape maintenance defects along the lateral cell cylinders.

### ReoM and ReoY affect the stability of MurA

Suppression of the Δ*gpsB* phenotype can be achieved by the accumulation of MurA (29). Consequently, MurA levels were determined in Δ*reoM* and Δ*reoY* mutant strains by Western blotting. MurA accumulated by at least eight-fold in comparison to the wild type in the absence of *reoM* or *reoY* (Fig. 2A), and reached similar levels to a mutant lacking *clpC*, which encodes the ATPase subunit of the ClpCP protease (Fig. 2A). MurAA, the *B. subtilis* MurA homologue, is degraded by the ClpCP protease *in vivo* (27). In order to test whether *reoM* and *reoY* exert their effect on MurA in a ClpC-dependent manner in *L. monocytogenes*, MurA levels were determined in Δ*clpC* Δ*reoM* and Δ*clpC* Δ*reoY* double mutants. The MurA levels in Δ*clpC*, Δ*reoM* and Δ*reoY* single mutants were the same as in Δ*clpC* Δ*reoM* and Δ*clpC* Δ*reoY* double mutant strains (Fig. 2B). Likewise, the MurA level in a mutant lacking *murZ*, previously shown to contribute to MurA stability (29), is not additive to the MurA level in Δ*clpC* cells (Fig. 2B). Therefore, ReoM, ReoY and MurZ likely affect the ClpCP-dependent degradation of MurA. Combinations of Δ*reoM,* Δ*reoY* and Δ*murZ* deletions did also not exert any additive effect on accumulation of MurA (Fig. S3A,B), further validating the conclusion that these genes all belong to the same pathway.

**Figure 2:**
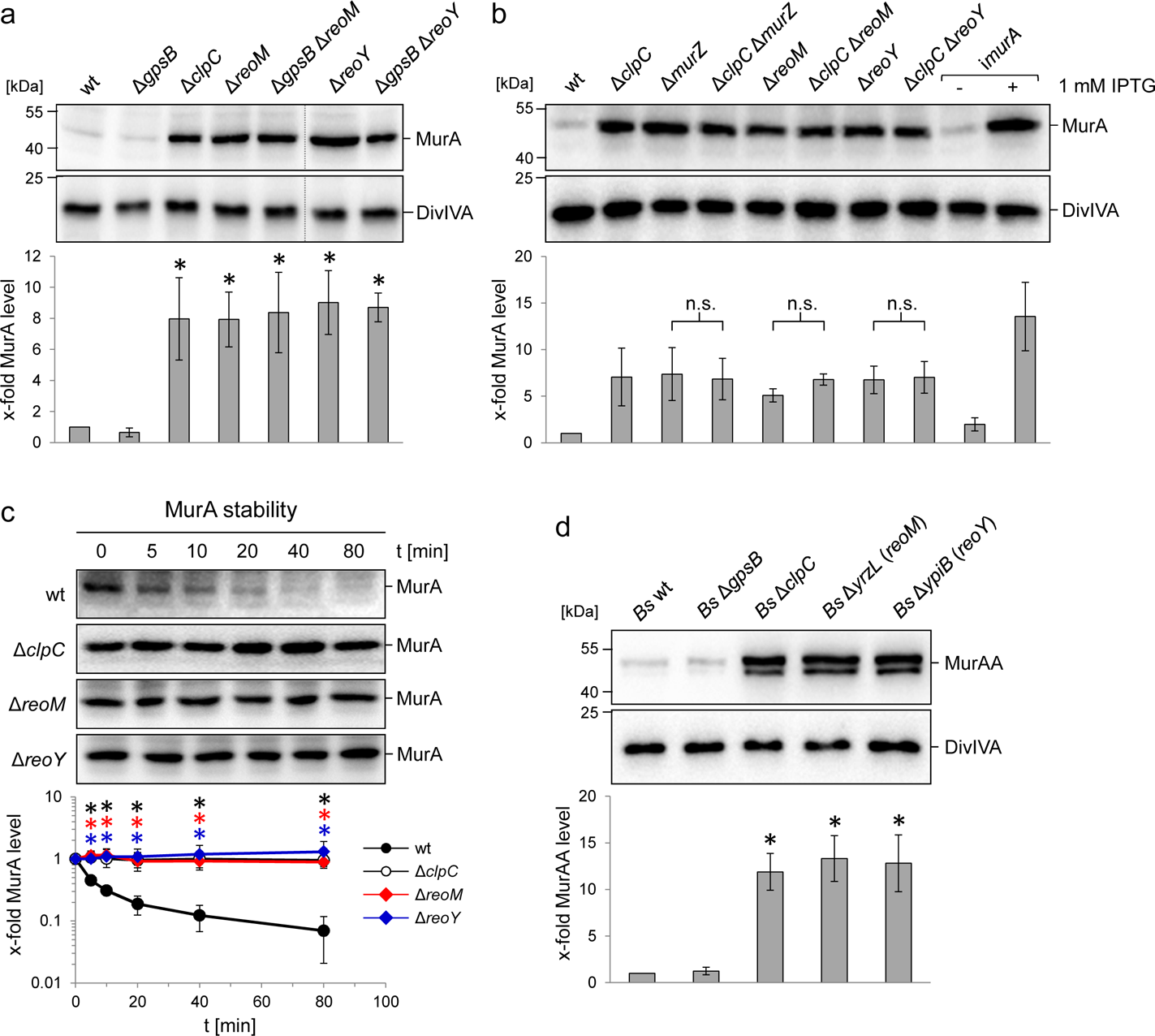
Effect of the *reoM, reoY* and *clpC* genes on levels of MurA in *L. monocytogenes* and MurAA in *B. subtilis*. (A) Effect of *reoM* and *reoY* deletions (single or when combined with *gpsB* deletion) on MurA (above) and DivIVA levels (middle) in *L. monocytogenes* strains EGD-e (wt), LMJR19 (Δ*gpsB*), LMSW30 (Δ*reoM*), LMSW32 (Δ*reoY*), LMJR137 (Δ*gpsB* Δ*reoM*) and LMJR120 (Δ*gpsB* Δ*reoY*) and quantification of MurA levels (below). Strain LMJR138 (Δ*clpC*) was included for comparison. Non-relevant lanes were excised from the blots (dotted lines). Average values ± standard deviations were shown (n=3). Statistically significant differences compared to wild type are marked by asterisks (*P*<0.05, *t*-test). (B) Effect of *reoM*, *reoY* and *murZ* deletions when combined with *clpC* deletion on MurA (above) and DivIVA levels (middle) in *L. monocytogenes* strains EGD-e (wt), LMJR138 (Δ*clpC*), LMJR104 (Δ*murZ*), LMJR171 (Δ*clpC* Δ*murZ*), LMSW30 (Δ*reoM*), LMSW50 (Δ*clpC* Δ*reoM*), LMSW32 (Δ*reoY*) and LMSW51 (Δ*clpC* Δ*reoY*) and quantification of MurA levels (below). Strain LMJR123 (I*murA*) grown in the presence or absence of IPTG was included for comparison. Average values and standard deviations were shown (n=3) and n. s. means not significant (*P*<0.05, *t*-test). (C) Western blots following MurA degradation *in vivo*. *L. monocytogenes* strains EGD-e (wt), LMJR138 (Δ*clpC*), LMSW30 (Δ*reoM*) and LMSW32 (Δ*reoY*) were grown to an OD_600_ of 1.0 and 100 µg/ml chloramphenicol was added. Samples were taken before chloramphenicol addition and after several time intervals to analyze MurA levels. MurA signals were quantified by densitometry and average values and standard deviations are shown (n=3). Statistically significant differences are marked with asterisks (*P*<0.05, *t*-test). (D) Effect of the *reoM* and *reoY* homologs *yrzL* and *ypiB*, respectively, on MurAA (above) and DivIVA levels (middle) of *B. subtilis* and quantification of MurAA levels (below). Strains BKE00860 (Δ*clpC*), BKE22180 (Δ*gpsB*), BKE22580 (Δ*ypiB*/*reoY*) and BKE27400 (Δ*yrzL*/*reoM*) were grown to mid-logarithmic growth phase before total cellular proteins were isolated. *B. subtilis* 168 (wt) was included as control. That MurAA is detected in two isoforms had been observed earlier but the reasons for this are not known (27). Average values and standard deviations were shown (n=3). Asterisks indicate statistically significant differences compared to wild type (*P*<0.05, *t*-test).

We then tested the hypothesis that ReoM and ReoY control proteolytic stability of MurA and followed MurA and DivIVA degradation over time in cells that had been treated with chloramphenicol to block protein biosynthesis. MurA was almost completely degraded in wild type cells 80 min after chloramphenicol treatment (Fig. 2C), whereas DivIVA was stable (Fig. S4). By contrast, no MurA degradation was observed in mutants lacking *clpC*, *reoM* or *reoY* (Fig. 2C), which together demonstrates that ReoM and ReoY are as important for MurA degradation as is ClpC.

### The effect of ReoM and ReoY on MurA levels is conserved

Homologues of the 90-residue ReoM protein are found across the entire Firmicute phylum, and include IreB, a substrate of the protein serine/threonine kinase IreK and its cognate phosphatase IreP from *Enterococcus faecalis* (38), whereas ReoY homologues are present only in the *Bacilli* and a *reoY* homologue has been identified as a Δ*ireK* suppressor in *E. faecalis* (39), but the function of both *E. faecalis* remains unkown. In *B. subtilis*, ReoM corresponds to YrzL (e-value 3e^-29^) and ReoY to YpiB (4e^-61^), but neither protein has been studied thus far. To assess whether YrzL and YpiB were also crucial for control of MurAA levels in *B. subtilis*, cellular protein extracts from *B. subtilis* Δ*yrzL* and Δ*ypiB* mutants were probed by Western blot (Fig. 2D). MurAA accumulated by at least 12-fold in these strains in comparison to the wild type. Furthermore, the amount of MurAA was also increased by 12-fold in the Δ*clpC* mutant. Taken together, these data indicate that ReoM and ReoY functions are conserved in both species. We thus propose to rename *lmo1503* (*yrzL*) as *reoM* (<u>re</u>gulator <u>o</u>f <u>M</u>urA(A) degradation) and analogously *lmo1921* (*ypiB*) as *reoY*.

Several other ClpC substrates are known in *B. subtilis*, including the glutamine fructose-6-phosphate transaminase GlmS and the acetolactate synthase subunit IlvB (40). The levels of both proteins were also significantly increased in *B. subtilis* Δ*reoM* and Δ*reoY* mutants (Fig. S5), indicating that ReoM and ReoY are required for degradation of ClpC substrates in general.

### ReoM and ReoY contribute to PG biosynthesis

In order to test whether MurA accumulation affected PG production, we tested the effect of enhanced MurA levels on resistance of *L. monocytogenes* against the cephalosporine antibiotic ceftriaxone. Artificial overproduction of MurA in strain LMJR116, which carries an IPTG-inducible *murA* gene in addition to the native copy on the chromosome, lead to a 12-fold increase of ceftriaxone resistance, while MurA depletion lowered ceftriaxone resistance (Tab. 1.). MurA levels are thus directly correlated with PG production, presumably leading to stimulation or impairment of PG biosynthesis during overproduction and depletion, respectively. In good agreement with the overproduction of MurA, ceftriaxone resistance of the Δ*clpC* mutant increased to the same degree as when MurA was overproduced (Tab. 1). Ceftriaxone resistance of Δ*reoM*, Δ*reoY* and Δ*murZ* mutants increased two- to three-fold (Tab. 1); this intermediate resistance level is probably explained by the presence of functional ClpCP in these strains. Nevertheless, these observations are consistent with a function of ReoM, ReoY and MurZ as regulators of ClpCP-dependent MurA degradation. Taken together, these results show that modulation of MurA levels effectively controls PG biosynthesis and also demonstrate that ReoM, ReoY and MurZ play an important role in its regulation.

**Table 1:** Minimal inhibitory concentrations (MIC) of ceftriaxone Average values and standard deviations are calculated from three independent experiments.

| strain | genotype | IPTG | MIC ceftriaxone [µg/ml] |
| --- | --- | --- | --- |
| EGD-e | wt | - | 8±0 |
| LMJR116 | wt+*murA* | - | 16±0 |
|  |  | + | 96±0 |
| LMJR123 | *imurA* | - | 2±1 |
|  |  | + | 14±2 |
| LMJR138 | Δ*clpC* | - | 96±0 |
| LMSW30 | Δ*reoM* | - | 19±5 |
| LMSW32 | Δ*reoY* | - | 21±5 |
| LMJR104 | Δ*murZ* | - | 24±0 |
| LMSW76 | $\Delta$ *prkP prkA* * | - | 21±5 |
| LMSW84<sup>a</sup> | *iprkA* | - | 0.05±0 |
|  |  | + | 1.5±0 |

<sup>a</sup> The *iprkA* strain showed residual growth on BHI agar plates not containing IPTG, even though
it required IPTG for growth in BHI broth.

### Phosphorylation and dephosphorylation of ReoM by PrkA and PrkP i*n vitro*

PrkA (encoded by *lmo1820*) and PrkP (*lmo1821*) are the *L. monocytogenes* homologs of *E. faecalis* IreK and IreP, respectively. Consequently, the pairwise interactions and biochemical properties of ReoM, the PrkA kinase domain (PrkA-KD) and the cognate phosphatase PrkP were investigated. All isolated proteins electrophoresed as single species in non-denaturing PAGE (lanes 1, 2, Fig. 3A; lanes 1-4, Fig. 3B). When ReoM was incubated with PrkA-KD, in the absence of ATP, a slower migrating species was observed and the individual bands corresponding to ReoM and PrkA-KD disappeared indicating that the slower migrating species was a ReoM:PrkA-KD complex (lane 3, Fig. 3A). When ReoM was incubated with PrkA-KD and Mg/ATP under the same conditions, free PrkA-KD was observed but no bands equivalent to ReoM and the ReoM:PrkA-KD complex remained; instead a new species was present, migrating faster in the gel than ReoM (lane 4, Fig. 3A), which is likely to be phosphorylated ReoM (P-ReoM). Intact protein liquid chromatography-mass spectrometry (LC-MS) analysis of ReoM isolated from PrkA-KD after phosphorylation revealed the addition of 79.9 Da in comparison to the mass of ReoM (10671.5 Da), which corresponds to the formation of a singly-phosphorylated ReoM product of 10751.4 Da (Fig. 3C, Fig. S6). MS/MS spectra obtained during peptide mass fingerprinting were also consistent with one phosphorylation event per protein chain: the mass of one ReoM peptide, spanning residues Asp5 to Lys22 with mass of 2151.89 Da, was increased by 79.96 Da after incubation with PrkA-KD and Mg/ATP. Analysis of the *y-* and *b-* ions in the MS/MS fragmentation spectrum of this peptide was consistent only with Thr7 as the sole phosphosite in ReoM (Fig. 3D). Finally, mutation of Thr7 to alanine completely abrogated the phosphorylation of ReoM by PrkA-KD when analysed by LC-MS (Fig. S7).

**Figure 3:**
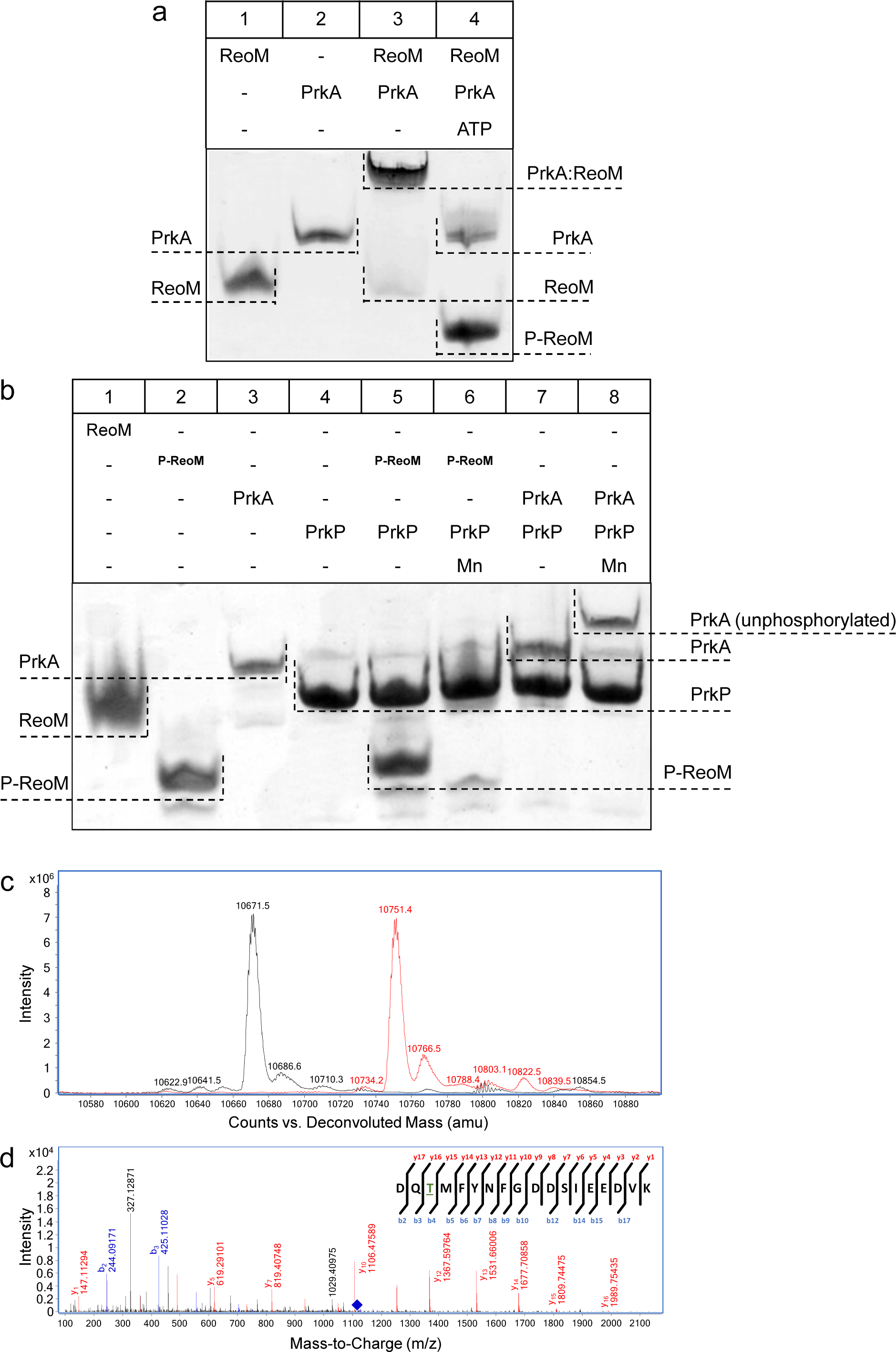
The PrkA/PrkP pair controls the phosphorylation status of ReoM. Non-denaturing, native PAGE analysis of the phosphorylation (A) and dephosphorylation (B) of ReoM *in vitro*. The components of each lane in the Coomassie-stained gel are annotated above the image and the position and identity of relevant bands is marked to the side. (C) LC-MS analysis of intact ReoM. The deconvoluted mass spectrum for non-phosphorylated ReoM (black) is overlaid over the equivalent spectrum for mono-phosphorylated ReoM, P-ReoM (red). (D) LC-MS/MS was used to perform peptide mapping analysis that revealed that Thr7 is the sole phosphosite of ReoM. The MS/MS fragmentation spectra of the phosphorylated peptide encompassing Asp5-Lys22 is presented with *b*-ion fragmentation in blue and *y*-ion fragmentation shown in red, whilst the precursor ion (m/z 1116.86, z=2+) is represented by a blue diamond.

The ability of PrkP, the partner phosphatase to PrkA in *L. monocytogenes,* to interact with and remove phosphoryl groups from PrkA-KD and P-ReoM was also tested *in vitro*. PrkA and purified P-ReoM were each incubated with PrkP in the absence and presence of MnCl_2_, since divalent cations are essential co-factors for the PPM phosphatase family to which PrkP belongs (41), and the products were analysed by non-denaturing PAGE. Unlike the situation with ReoM and PrkA-KD, no stable protein:protein complexes were formed either in the presence or absence of endogenous MnCl_2_ (Fig. 3B). The incubation of P-ReoM with PrkP and manganese resulted in the almost complete disappearance of the band corresponding to P-ReoM (lane 6, Fig. 3B) in comparison to the same reaction conducted without the addition of MnCl_2_ (lane 5, Fig. 3B). The new band, corresponding to ReoM alone in lane 6, is masked by that for PrkP which migrates similarly to ReoM (lanes 1 and 4, Fig. 3B) under these electrophoresis conditions. The presence of unphosphorylated ReoM and the absence of P-ReoM was confirmed by LC-MS (Fig. S8). When incubated with PrkP in the presence of manganese ions, the band for PrkA-KD electrophoresed more slowly than for PrkA-KD in isolation (lanes 3 and 8, Fig. 3B), indicating that PrkA-KD had been dephosphorylated by PrkP. LC-MS analysis of PrkA-KD that had been incubated with PrkP/MnCl_2_ yielded a single major species of 37,413.2 Da, consistent with the predicted mass of the expressed recombinant construct, and the absence of a peak corresponding to phosphorylated PrkA-KD, P-PrkA-KD (Fig. S9). Therefore, PrkA-KD is capable of autophosphorylation even when expressed in a heterologous host, consistent with previous observations made for similar PASTA-eSTKs from other Gram-positive bacteria (42, 43). Finally, in the absence of MnCl_2_ no change in electrophoretic mobility was observed for P-PrkA-KD (lane 7, Fig. 3B).

### Phosphorylation of ReoM at threonine 7 is essential for viability

PrkA phosphorylates ReoM on Thr7 and PrkP reverses this reaction *in vitro*; ReoM phosphorylation at Thr7 *in vivo* has also been observed by phosphoproteomics (44). In the absence of molecular details on the impact of Thr7 phosphorylation we determined the importance of this phosphorylation *in vivo* by engineering a phospho-ablative T7A exchange in an IPTG-inducible allele of *reoM* and introduced it into the Δ*reoM* mutant background. Deletion, depletion or expression of wildtype *reoM* had no effect on growth in strains LMSW30 (Δ*reoM*) and LMSW57 (i*reoM,* i - is used to denote IPTG-dependent alleles throughout the manuscript) at 37°C. Likewise, strain LMSW52 (i*reoM T7A*) grew normally in the absence of IPTG. However, the *reoM* mutant with the T7A mutation did not grow at all in the presence of IPTG, when expression of the phospho-ablative *reoM* T7A allele was induced (Fig. 4A), suggesting that phosphorylation of ReoM at Thr7 is essential for the viability of *L. monocytogenes*. Since ReoM influences the proteolytic stability of MurA, we determined the cellular amount of MurA in strains expressing the T7A variant of ReoM. For this purpose, strains LMSW57 (i*reoM*) and LMSW52 (i*reoM T7A*) were initially cultivated in plain BHI broth. At an OD_600_ of 0.2, IPTG was added to a final concentration of 1 mM and cells were harvested 2 hours later. Strain LMSW57 (i*reoM*) showed Δ*clpC*-like MurA accumulation (around seven-fold in this experiment) when cultured in the absence of IPTG, but MurA was present at wild type levels in the presence of IPTG (Fig. 4C). The strain with the T7A exchange also accumulated MurA to a Δ*clpC*-like extent in the absence of IPTG. However, around 10% of the wild type MurA levels could be detected in cells expressing the *reoM T7A* allele (Fig. 4C). These data demonstrate that Thr7 in ReoM is of special importance for the proteolytic stability of MurA. In agreement with these results, IPTG was toxic for the i*reoM T7A* mutant in a disc diffusion assay and rendered this strain hypersensitive to ceftriaxone (Fig. 4D).

**Figure 4:**
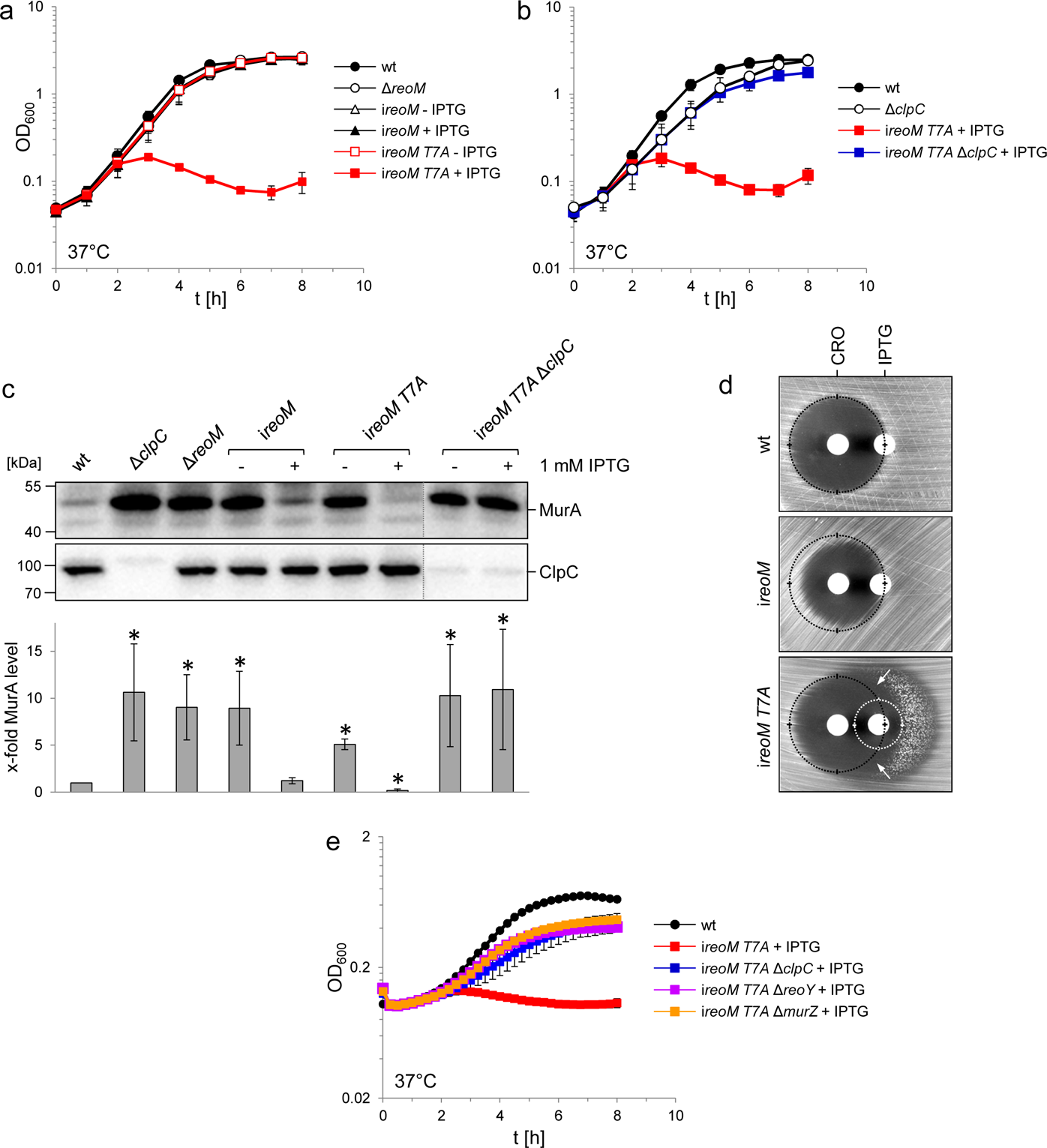
A ReoM T7A exchange affects growth and MurA levels in a ClpC-dependent manner. (A) Lethality of the *reoM T7A* mutation in *L. monocytogenes*. *L. monocytogenes* strains EGD-e (wt), LMSW30 (Δ*reoM*), LMSW57 (i*reoM*) and LMSW52 (i*reoM T7A*) were grown in BHI broth ± 1 mM IPTG at 37°C. The experiment was repeated three times and average values and standard deviations are shown. (B) Suppression of *reoM T7A* lethality by deletion of *clpC*. *L. monocytogenes* strains EGD-e (wt), LMJR138 (Δ*clpC*), LMSW52 (i*reoM T7A*) and LMSW72 (i*reoM T7A* Δ*clpC*) were grown in BHI broth ± 1 mM IPTG at 37°C. The experiment was repeated three times and average values and standard deviations are shown. (C) Western blot showing cellular levels of MurA (top) and ClpC (middle) in the strains included in panels A and B. For this experiment, strains were grown in BHI broth not containing IPTG at 37°C. IPTG (1 mM) was added to the cultures at an OD_600_ of 0.2 and the cells were collected 2 hours later. Irrelevant lanes were removed from both blots (dotted lines). Quantification of MurA signals by densitometry is shown below the Western blots. Average values and standard deviations calculated from three independent experiments are shown. Asterisks indicate statistically significant differences (*P*<0.05, *t*-test). (D) ReoMT7A expression sensitizes *L. monocytogenes* against ceftriaxone. Synergism between ceftriaxone and IPTG in the i*reoMT7A* strain LMSW52 in a disc diffusion assay with filter discs containing 50 mg/ml ceftriaxone (CRO, left) and 1 mM IPTG (right). For comparison, wild type levels of growth inhibition by ceftriaxone are marked with black circles. Zone of growth inhibition by IPTG in the i*reoM T7A* mutant is marked with a white circle. Please note that strain LMSW52 shows hetero-resistance against IPTG (two zones of growth inhibition with different resistance levels). Arrows mark the zones of synergism between ceftriaxone and IPTG. (E) Contribution of ReoY and MurZ to the lethal *reoM T7A* phenotype. *L. monocytogenes* strains EGD-e (wt), LMSW52 (i*reoM T7A*), LMSW72 (i*reoM T7A* Δ*clpC*), LMSW123 (i*reoM T7A* Δ*reoY*) and LMSW124 (i*reoM T7A* Δ*murZ*) were grown in BHI broth containing 1 mM IPTG and growth at 37°C was recorded in a microplate reader. Average values and standard deviations were calculated from an experiment performed in triplicate.

### Lethality of the *reoM T7A* mutations depends on ClpC

That MurA is rapidly degraded in cells expressing *reoM T7A* implies that phosphorylation/dephosphorylation of ReoM at Thr7 controls ClpCP-dependent MurA degradation. MurA is an essential enzyme in *L. monocytogenes* (29), and stimulation of ClpCP-dependent MurA degradation in the *reoM T7A* mutant would provide an explanation for the lethality of this mutation. In order to address this possibility, we deleted *clpC* in the conditional i*reoM T7A* background. This strain grew even in the presence of IPTG, a compelling demonstration that the removal of *clpC* suppressed the lethality of the *reoM T7A* mutation (Fig. 4B). MurA also accumulated to the same degree as in the Δ*clpC* mutant in this strain (Fig. 4C), suggesting that inactivation of the ClpCP-dependent degradation of MurA overcame the lethal effect of the T7 mutation in *reoM* and this suggests that ClpCP acts downstream of ReoM. We next wondered whether deletion of *reoY* and *murZ* would have a similar effect and deleted these genes in the *reoM T7A* mutant. As can be seen in Fig. 4E, deletion of either gene overcame the lethality of *reoM T7A*, indicating that ReoY and MurZ must also act downstream of ReoM.

### Crystal structure of ReoM, a homologue of *Enterococcus faecalis* IreB

Purified ReoM yielded crystals that diffracted to a maximum resolution of 1.6 Å. The NMR structure of IreB (PDBid 5US5) (45) was used to solve the structure of ReoM by molecular replacement (Fig. 5A). The data collection and refinement statistics for the ReoM structure are summarised in Tab. 2. ReoM shares the same overall fold as IreB (45), each containing a compact 5-helical bundle (4 standard α-helices and one single-turned 3_10_-helix between residues 52 and 54) with short loops between the secondary structure elements, which are defined above the sequence alignment in Fig. 5B. Other than IreB (45), there are no structural homologues of ReoM with functional significance in the PDB. The helical bundles in both ReoM and IreB associate into homodimers with α-helices two and four from each protomer forming the majority of the homodimer interface (Fig. 5A), and these residues are highlighted in Fig. 5B. In agreement with the IreB structural analysis, 1200 Å^2^ of surface area is buried in the ReoM dimer interface, representing 9% of the total solvent accessible surface area. The similarity of the monomers and the dimeric assemblies of ReoM and IreB is underlined by the 1.5 and 1.7 Å r.m.s.d. values, respectively, on global secondary structure superposition matching 74 Cα atoms from each protomer in the comparison.

**Figure 5:**
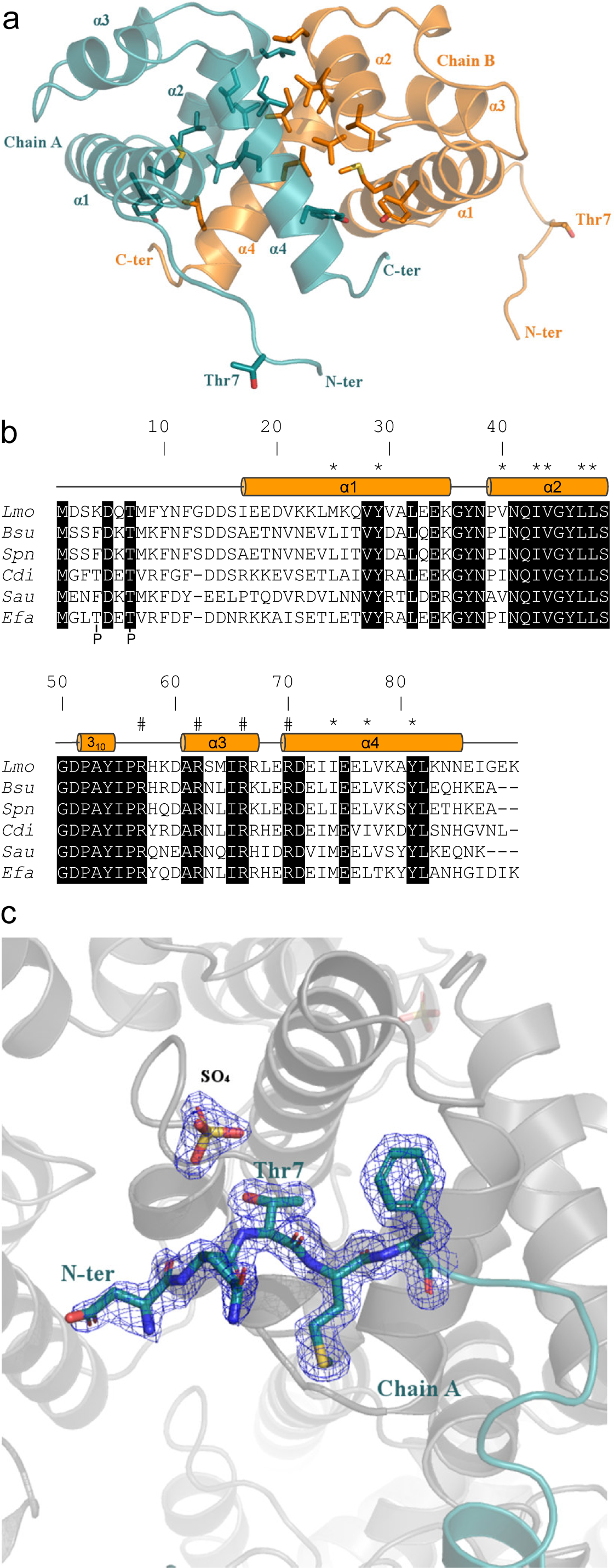
Crystal structure of ReoM. (A) The structure of ReoM depicted as a cartoon with each protomer in the dimer coloured separately (cyan and orange). The secondary structure elements are numbered according to their position in the amino acid sequence. Thr7 and some of the key amino acids in the dimer interface and the hydrophobic core are drawn as stick figures. (B) Sequence alignment of ReoM (*Lmo*) and its homologues from *Bacillus subtilis* (*Bsu*), *Streptococcus pneumoniae* (*Spn*), *Clostridium difficile* (*Cdi*) and *Staphylococcus aureus* (*Sau*) with the sequence of IreB from *Enterococcus faecalis* (*Efa*) underneath. Amino acid sequence numbers pertain to ReoM and the site of phosphorylation in ReoM (Thr7) and the twin phosphorylations in IreB (minor site: Thr4; major site: Thr7) are highlighted. Invariant amino acids are shaded black, residues in the ReoM dimer interface have an asterisk above, and the secondary structure elements are defined by cylinders above the alignment. Arginine residues mutated in this study are indicated by a hashtag above the alignment. (C) The final 2F_obs_-F_calc_ electron density map, contoured at a level of 0.42 e^-^/Å^3^, of the N-terminal region in the immediate vicinity of Thr7 in chain A of the ReoM dimer indicates that the protein model could be built with confidence even though this region contains no secondary structure elements.

**Table 2:** Summary of the data collection and refinement statistics for ReoM.

| Data collection |  |
| --- | --- |
| Beamline | Diamond I03 |
| Wavelength (Å) | 0.976 |
| Resolution (Å) | 74.45-1.60 (1.63-1.60)* |
| Space group | P 2<sub>1</sub> 2<sub>1</sub> 2<sub>1</sub> |
| *a*, *b*, *c* (Å) | 38.79, 58.62, 74.45 |
| $\alpha$, $\beta$, $\gamma$ (°) | 90, 90, 90 |
| R<sub>pim</sub> | 0.064 (0.533) |
| CC (1/2) (%) | 98.6 (62.0) |
| <I>/<$\sigma$(I)> | 8.2 (2.2) |
| Completeness (%) | 99.8 (99.8) |
| Redundancy | 4.8 (4.9) |
| Total observations | 111229 (5581) |
| Unique reflections | 23059 (1129) |
| Refinement |  |
| R<sub>work</sub> (%) | 15.3 |
| R<sub>free</sub> (%) | 21.4 |
| Solvent content (%) | 38.0 |
| # atoms |  |
| Protein | 1399 |
| Ligand/ion | 20 |
| Water | 94 |
| B-factors (Å<sup>2</sup>) |  |
| Protein | 26.4 |
| Ligand/ion | 50.5 |
| Water | 37.7 |
| R.m.s deviations |  |
| Bonds (Å) | 0.015 |
| Angles (°) | 1.79 |

\*Where values in parentheses refer to the highest resolution shell.

Other than the compact helical bundle of ReoM, there is a ∼16 residue-long N-terminal tail, with B-factors 25% higher than the rest of the protein, prior to the start of α-helix 1 at residue Ile17. The equivalent N-terminal region is also disordered in the NMR structure of IreB (45). Despite the absence of secondary structure, the ReoM model covering this region could be built with confidence from Asp5 in chain A and Asp2 in chain B (Fig. 5C). Consequently, it is possible to visualise Thr7, the target for phosphorylation by PrkA, in the flexible N-terminal region in both chains. The side chain of Thr7 in both chains makes no intramolecular interactions and is thus amenable to phosphorylation by PrkA. The extended N-terminal regions are at least partially stabilised by crystal lattice interactions that, in chain B, include Phe9 and Tyr10 forming a network of hydrophobic interactions with other aromatic residues from symmetry equivalent molecules, including contributions from another copy of Phe9 and Tyr10. Phosphorylation could force a change in oligomeric state, as observed quite commonly in response regulators in order to bind more effectively to promoter regions to effect transcription (46). However, analysis of the oligomeric state of P-ReoM by size exclusion chromatography revealed that the protein behaved in solution the same as to unphosphorylated ReoM (Fig. S10).

Unlike the packing arrangement of Thr7 in chain B, the local symmetry surrounding Thr7 in chain A might provide some information of potential functional significance to the phosphorylated form of ReoM. Here, a sulphate ion (a component of the crystallisation reagent) is hydrogen bonded to the sidechain of Thr7 and hence mimics, to some degree, what the phosphorylated protein may look like (Fig. 5C). The sulphate ion is captured by a positively-charged micro-environment incorporating residues Lys35, Arg57, His58 and Arg62 from a symmetry-equivalent molecule. ReoM could react to phosphorylation on Thr7 by a substantial movement of the N-terminal tail to interact with conserved, positively-charged amino acids on the protein surface. We identified a single cluster of arginines (Arg57 [57% conserved], Arg62 [99%], Arg66 [76%], Arg70 [98%]) in close spatial proximity with levels of conservation amongst all 2909 ReoM homologues present at NCBI approaching that of Thr7 (96%,) and replaced them by alanines. Whereas the R66A and R70A mutations were without any effect on growth (data not shown), expression of ReoM R57A and R62A mutations were as lethal as expression of ReoM T7A (Fig. S11). Thus, Arg57 and Arg62 might co-ordinate P-Thr7, stabilising the conformation and position of the flexible N-terminal region (Fig. S12). Despite multiple attempts, however, no crystals of P-ReoM could be grown and the molecular consequences of ReoM phosphorylation remain to be determined.

### Control of MurA stability and PG biosynthesis by the PrkA/PrkP protein kinase/phosphatase pair

To study the contribution of the PrkA/PrkP couple to PG biosynthesis in more detail, we aimed to construct *prkA* and *prkP* deletion mutants, but failed to delete *prkA*. However, *prkA* could be deleted in the presence of an IPTG-inducible ectopic *prkA* copy and the resulting strain (LMSW84) required IPTG for growth (Fig. 6A), demonstrating the essentiality of this gene. The essentiality of *prkA* in our hands is consistent with results by others who have also shown that *prkA* can only be inactivated in the presence of a second copy (47). Repeated attempts to delete *prkP* finally yielded a single Δ*prkP* clone (LMSW76). Genomic sequencing of this strain, which grew at a similar rate to wild type (Fig. 6A), confirmed the successful deletion of *prkP* but also identified a trinucleotide deletion in the *prkA* gene (designated *prkA*\*), effectively removing the complete codon of Gly18 that is part of a conserved glycine-rich loop important for ATP binding (48). Presumably, this mutation reduces the PrkA kinase activity to balance the absence of PrkP. By contrast, *prkP* could be deleted readily in the presence of a second IPTG-dependent copy of *prkP* and growth of the resulting strain (LMSW83) did not require IPTG, most likely explained by promoter leakiness in the absence of IPTG (Fig. 6A). The viability of the i*prkP* mutant shows that the *prkP* deletion had no polar effects on the expression of the downstream *prkA*. That *prkA* and *prkP* are both essential suggests that some of their substrates must be phosphorylated and unphosphorylated, respectively, to be active. Next, the effect of *prkA* and *prkP* mutations on MurA accumulation was analyzed by Western blotting. Intermediate MurA accumulation was evident in the Δ*prkP prkA\** strain, while full accumulation of MurA was observed in PrkP-depleted cells. By contrast, no MurA was detected in cells depleted for PrkA (Fig. 6B). Therefore, PrkA and PrkP inversely contribute to the accumulation of MurA, suggesting that phosphorylated ReoM favors MurA accumulation, while un-phosphorylated ReoM counteracts this process. In good agreement, depletion of PrkA strongly increased ceftriaxone susceptibility, while inactivation of *prkP* caused increased ceftriaxone resistance (Tab. 1).

**Figure 6:**
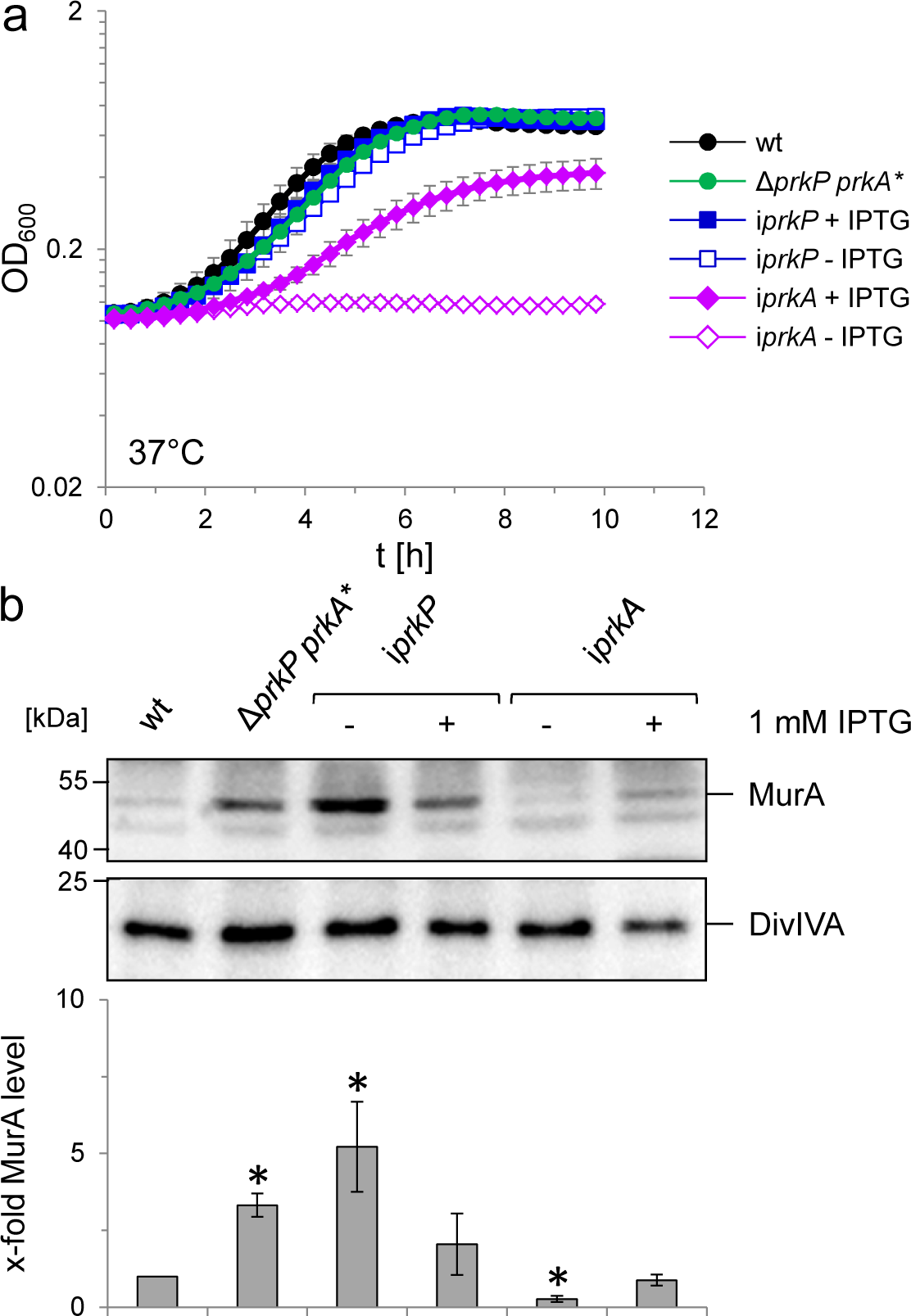
Effect of *prkA* and *prkP* mutations on growth and MurA levels of *L. monocytogenes*. (A) Contribution of PrkA and PrkP to *L. monocytogenes* growth. *L. monocytogenes* strains EGD-e (wt), LMSW76 (Δ*prkP prkA\**), LMSW83 (i*prkP*) and LMSW84 (i*prkA*) were grown in BHI broth containing or not containing 1 mM IPTG at 37°C in a microtiter plate reader. The experiment was repeated three times and average values and standard deviations are shown. (B) Contribution of PrkA and PrkP to MurA stability. Western blots showing cellular levels of MurA (top) and DivIVA (middle) in the same set of strains as in panel A and quantification of MurA signals by densitometry (below). Average values and standard deviations calculated from three independent experiments are shown. Asterisks indicate statistically significant differences (*P*<0.05, *t*-test).

### Deletion of *reoM*, *reoY* or *clpC* eliminates *prkA* essentiality

In order to test whether the essentiality of *prkA* could be explained by stimulated MurA degradation through ClpCP, we first tested the effect of *clpC* on the essentiality of *prkA*. For this purpose, *clpC* was removed from the i*prkA* strain and growth of the resulting strain (LMSW91) was tested. In contrast to the parental i*prkA* strain (LMSW84), which required IPTG for growth, strain LMSW91 was viable without IPTG (Fig. 7A) thus confirming that the essentiality of PrkA depends on ClpC. We next wondered whether ReoM and ReoY were also required for PrkA essentiality and consequently deleted their genes from the i*prkA* background to test this. Again, the resulting strains did not require IPTG for growth in contrast to the parental i*prkA* strain (Fig. 7A). In good agreement with these findings, deletion of *clpC*, *reoM* or *reoY* all stabilized MurA in PrkA-depleted cells (Fig. 7B), showing that the stimulated degradation of MurA that we observe in cells depleted for PrkA (Fig. 6B) is dependent on any one of these three proteins. These results together permit a model of genetic interactions to be proposed (Fig. 7C) that starts with PrkA and its downstream substrate ReoM. ReoY, MurZ and ClpC in turn are positioned downstream of ReoM (as indicated by the experiments shown in Fig. 4D) to control MurA stability. To further substantiate this concept, physical interactions between ReoM, ReoY, ClpC, ClpP and MurA were determined in bacterial two hybrid experiments, which revealed that ReoY interacted with ClpC, ClpP and ReoM. In turn, ReoM interacted with MurA (Fig. 7C, Fig. S13), which suggests that ReoM and ReoY could bridge the interaction of ClpCP with its substrate MurA.

**Fig. 7:**
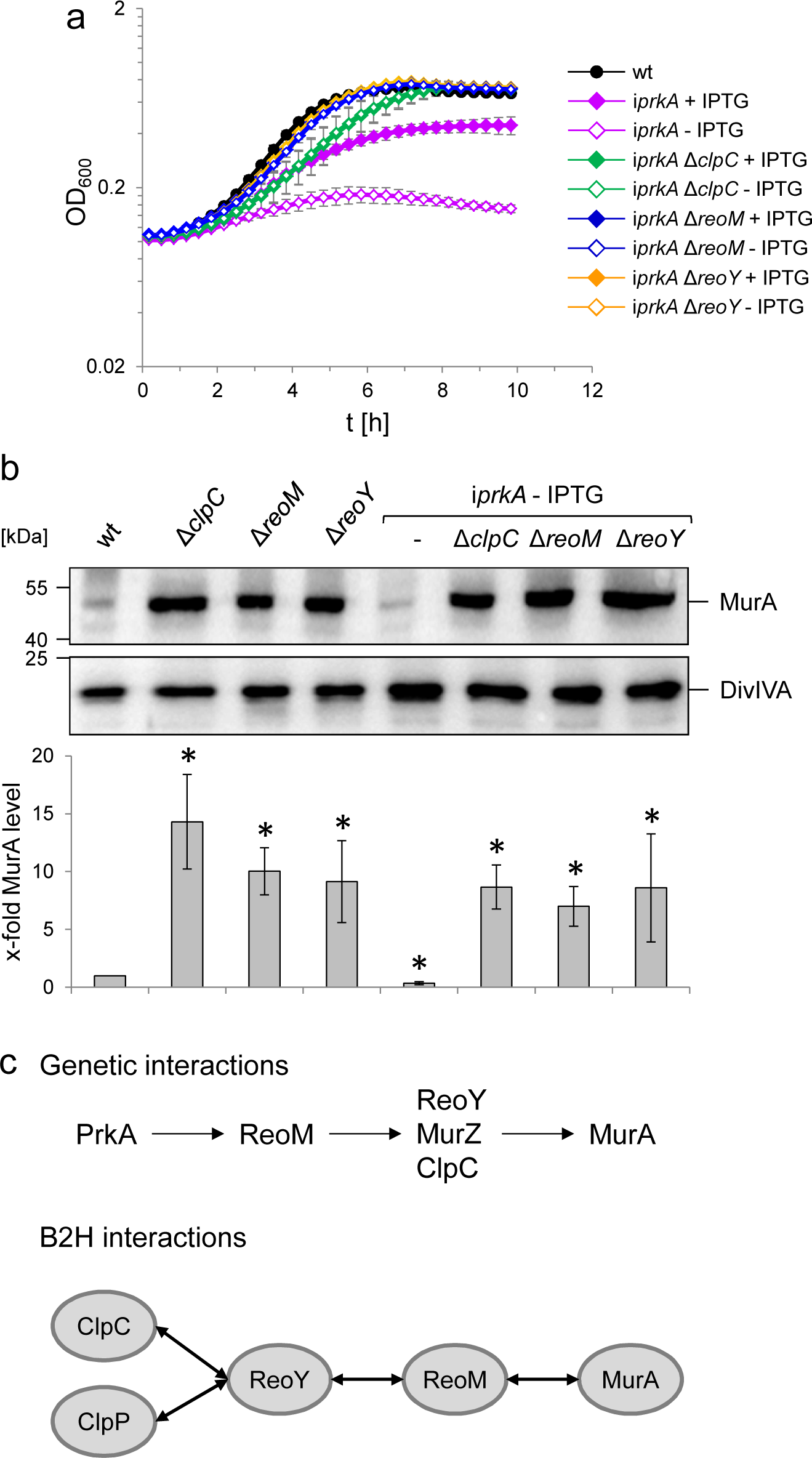
PrkA essentiality depends on *reoM*, *reoY* and *clpC*. (A) Effect of *reoM*, *reoY* and *clpC* deletions on *prkA* essentiality. *L. monocytogenes* strains EGD-e (wt), LMSW84 (i*prkA*), LMSW89 (i*prkA* Δ*reoM*), LMSW90 (i*prkA* Δ*reoY*) and LMSW91 (i*prkA* Δ*clpC*) were grown in BHI broth ± 1 mM IPTG at 37°C in a microtiter plate reader. The experiment was repeated three times and average values and standard deviations are shown. (B) *clpC, reoM* and *reoY* deletions overcome MurA degradation in PrkA-depleted cells. Western blot showing MurA levels in *L. monocytogenes* strains EGD-e (wt), LMJR138 (Δ*clpC*), LMSW30 (Δ*reoM*), LMSW32 (Δ*reoY*), LMSW84 (i*prkA*), LMSW89 (i*prkA* Δ*reoM*), LMSW90 (i*prkA* Δ*reoY*) and LMSW91 (i*prkA* Δ*clpC*, top). PrkA wild type strains were grown in BHI broth at 37°C to mid-exponential growth phase before protein isolation. A parallel DivIVA Western blot was used as loading control (middle). Quantification of MurA signals by densitometry (below). Average values and standard deviations calculated from three independent experiments are shown. Asterisks indicate statistically significant differences (*P*<0.05, *t*-test). (C) Illustrations summarizing the genetic interactions detected in this work between PrkA, ReoM, ReoY, MurZ, ClpC and MurA (above) and the physical interactions between them as detected by bacterial two hybrid analysis (bottom, raw data are shown in Fig. S13).

## DISCUSSION

With ReoM we have identified a missing link in a regulatory pathway that enables Firmicute bacteria to activate PG biosynthesis under conditions damaging their cell walls. In *L. monocytogenes*, the sensory module of this pathway comprises the membrane integral protein kinase PrkA and the cognate protein phosphatase PrkP, their newly discovered substrate ReoM and the associated factors ReoY and MurZ, which together regulate ClpCP activity, the effector protease that acts on MurA (Fig. 8). It has been demonstrated previously that the kinase activity of PrkA homologues was activated by muropeptides (17, 49) or the PG precursor lipid II (18). Muropeptides were released from the cell wall during normal PG turnover, and their release was intensified when PG hydrolysis prevailed over PG biosynthesis (10, 50), whereas blocking PG chain elongation by moenomycin treatment caused the accumulation of lipid-linked PG precursors (51). Thus, both types of molecules accumulated when PG biosynthesis was inhibited and could represent useful signals for detecting cell wall-damaging situations. Our data are consistent with a model in which PrkA-phosphorylated ReoM no longer activates ClpCP, which leads to MurA stabilization and the activation of PG biosynthesis (Fig. 8). In *B. subtilis*, this effect is supported by stabilization of GlmS (Fig. S4A), another ClpCP substrate but which acts in front of MurA as the first enzyme of the UDP-Glc*N*Ac-generating GlmSMU pathway.

**Fig. 8:**
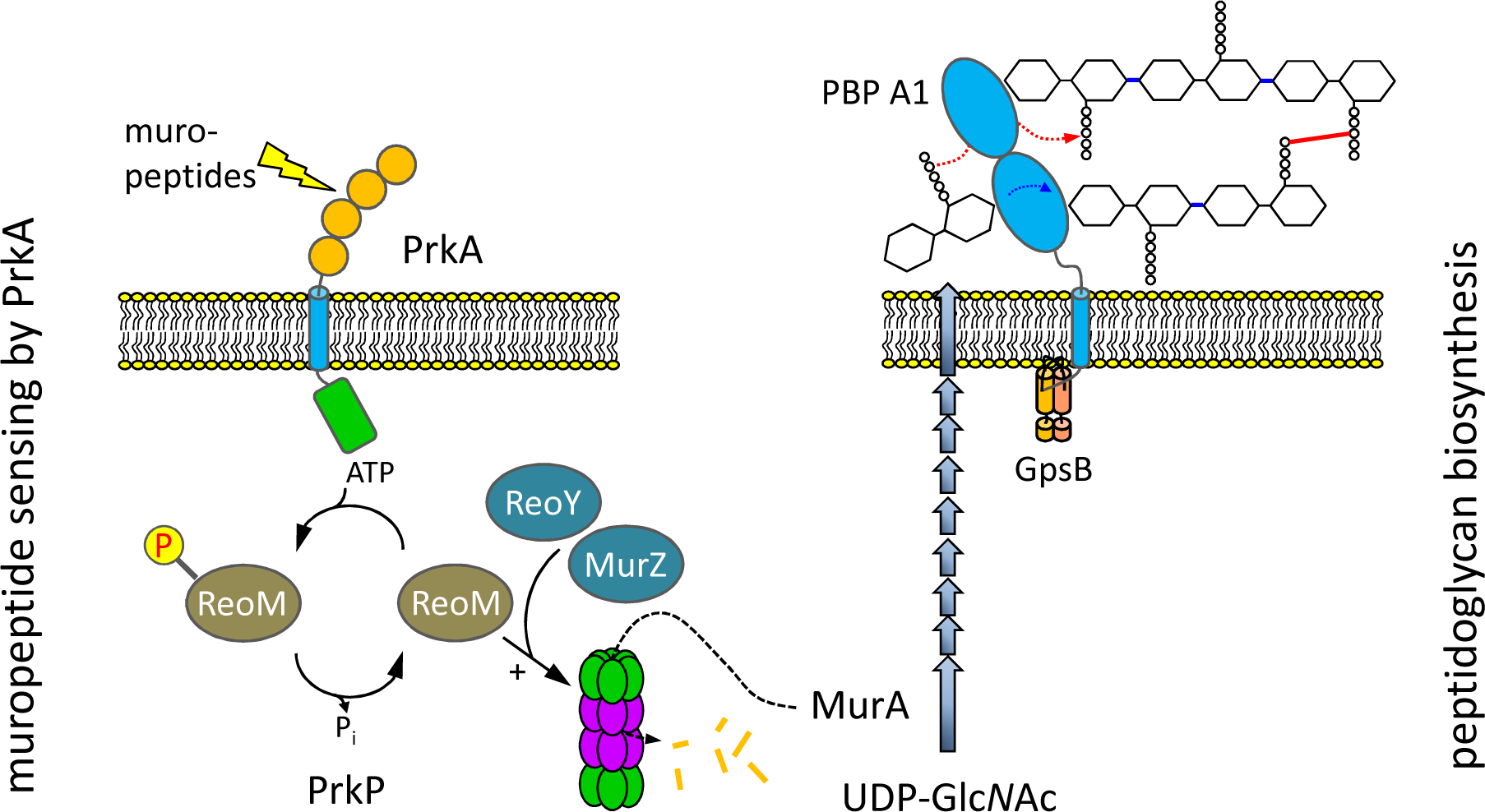
ReoM links PrkA-dependent muropeptide sensing with peptidoglycan biosynthesis. Model illustrating the role of ReoM as substrate of PrkA and as regulator of ClpCP. Cell wall damage is sensed by PrkA through recognition of free muropeptides upon which PrkA phosphorylates ReoM. In its unphosphorlyated form, ReoM is an activator of ClpCP-dependent degradation of MurA, the first enzyme of peptidoglycan biosynthesis, and ReoY and MurZ contribute to this process. By phosphorylating ReoM, PrkA prevents ClpCP-dependent MurA degradation so that MurA accumulates and peptidoglycan biosynthesis can occur. Please note that there is a lesser degree of conservation in the fourth PASTA domain of PrkA.

How ReoM and ReoY exert their effect on ClpCP is currently unknown, but a fascinating possibility would be a function like to that of an adaptor protein to target protein substrates to ClpCP for degradation. Several ClpC adaptors for different substrates are known in *B. subtilis* (52, 53), but an adaptor for *Bs*MurAA is not among them (27, 52). Like ReoM, the ClpC adaptor McsB from *B. subtilis* is also subject to phosphorylation, but - unlike ReoM - it targets its substrate CtsR to the ClpCP machinery only when phosphorylated (54). Either ReoM or ReoY could act as this adaptor, leaving a subsidiary function for the other respective protein. Alternatively, both proteins could work in tandem, where each of them is equally needed for ClpCP-dependent MurA degradation since the phenotypes of *reoM* and *reoY* mutants were identical with respect to MurA stability. However, overexpression or deletion of *reoM* altered the phenotype of the Δ*gpsB* mutant, but that of *reoY* was without phenotype (Fig. S1, Fig. S2). ReoY, restricted to the *Bacilli*, also showed a narrower phylogenetic distribution than ReoM, which is found across different Firmicutes (Fig. 5B). Thus, it seems that ReoM might have a more generalized role, whereas ReoY could play a subordinate function in control of MurA degradation by ClpCP. The role of the MurA homologue MurZ in this process is entirely unclear, but our genetic data now place it downstream of ReoM (Fig. 7C). Furthermore, arginine phosphorylation targets proteins to ClpCP for degradation (55). *L. monocytogenes* MurA contains 17 arginines and MurAA of *B. subtilis* has been found in complex with the protein arginine phosphatase YwlE (56). The possibility that MurA proteins could also require arginine phosphorylation to be accepted as a substrate by ClpCP offers additional control possibilities for ReoM/ReoY/MurZ to modulate MurA levels.

A screen for *gpsB* suppressors in *S. pneumoniae* did not yield *reoM* mutations (and these strains do not contain *reoY*, consistent with a subordinate function for this gene), but instead suppressor mutations were found that affect *phpP*, which encodes a Ser/Thr protein phosphatase that acts in concert with StkP, the PASTA-eSTK of this organism (57, 58). Absence or inactivation of PhpP triggered an increase in StkP-dependent protein phosphorylation levels in the pneumococcus (57, 59). It is tempting to speculate that loss of PhpP activity in this suppressor also triggers P-ReoM formation that, according to our model, would help to stabilize MurA and thus suppress the Δ*gpsB* phenotype. Interestingly, another *S. pneumoniae gpsB* suppressor was identified that carries a duplication of a ∼150 kb genomic fragment (57), a region that includes the open reading frame for MurA. Suppression of the *gpsB* phenotype in this instance could also work via MurA accumulation, but this time due to a gene dosage effect.

It is becoming increasingly evident that control of PG biosynthesis in response to cell wall derived signals, via PASTA-eSTKs, is a regulatory capacity common to Firmicutes and Actinobacteria. CwlM is the critical kinase substrate in the actinobacterium *M. tuberculosis* that, when phosphorylated by PknB, binds to and activates MurA (24). Homologues of CwlM are not present in *L. monocytogenes* or *B. subtilis* and instead these bacteria adjust their MurA levels by controlling MurA turnover in response to PrkA-dependent phosphorylation of ReoM. Consequently, both mechanisms activate PG biosynthesis in a PrkA-dependent manner either by activation or stabilization of MurA. Presumably *B. subtilis*, and other endospore forming bacteria, re-start PG biosynthesis at the onset of germination in a similar way. Germination of *B. subtilis* spores can be triggered by muropeptides in a manner that depends upon PrkC (49), the PASTA-eSTK of *B. subtilis* (60). Even though *Bs*PrkC phosphorylates multiple substrates (61), whose individual contribution to germination is not known precisely, phosphorylation of ReoM (*aka* YrzL) could be required to restart PG biosynthesis in germinating *B. subtilis* cells by stabilizing MurAA. Moreover, an *E. faecalis* mutant lacking the PASTA-eSTK IreK was more susceptible to ceftriaxone but overexpression of *Ef*MurAA overcame this defect (62). This implies the possibility that unphosphorylated IreB together with the ReoY homologue of this organism, OG1RF_11272 (39), might stimulate MurAA proteolysis in *E. faecalis* as well. Taken together it seems that observations made in different Firmicutes are in good agreement with the PrkA→ReoM/ReoY→ClpC→MurA signaling sequence that we propose. The open questions that remain on the molecular mechanism of ClpCP control by ReoM and ReoY will be addressed by future experiments.

## MATERIALS AND METHODS

### Bacterial strains and growth conditions

Tab. 3 lists all strains used in this study. Strains of *L. monocytogenes* were cultivated in BHI broth or on BHI agar plates. *B. subtilis* strains were grown in LB broth at 37°C. Antibiotics and supplements were added when required at the following concentrations: erythromycin (5 µg/ml), kanamycin (50 µg/ml), X-Gal (100 µg/ml) and IPTG (as indicated). *Escherichia coli* TOP10 was used as host for all cloning procedures (63). Minimal inhibitory concentrations against ceftriaxone were determined as described previously (64) using E-test strips with a ceftriaxone concentration range of 0.016 - 256 µg/ml.

**Table 3:**
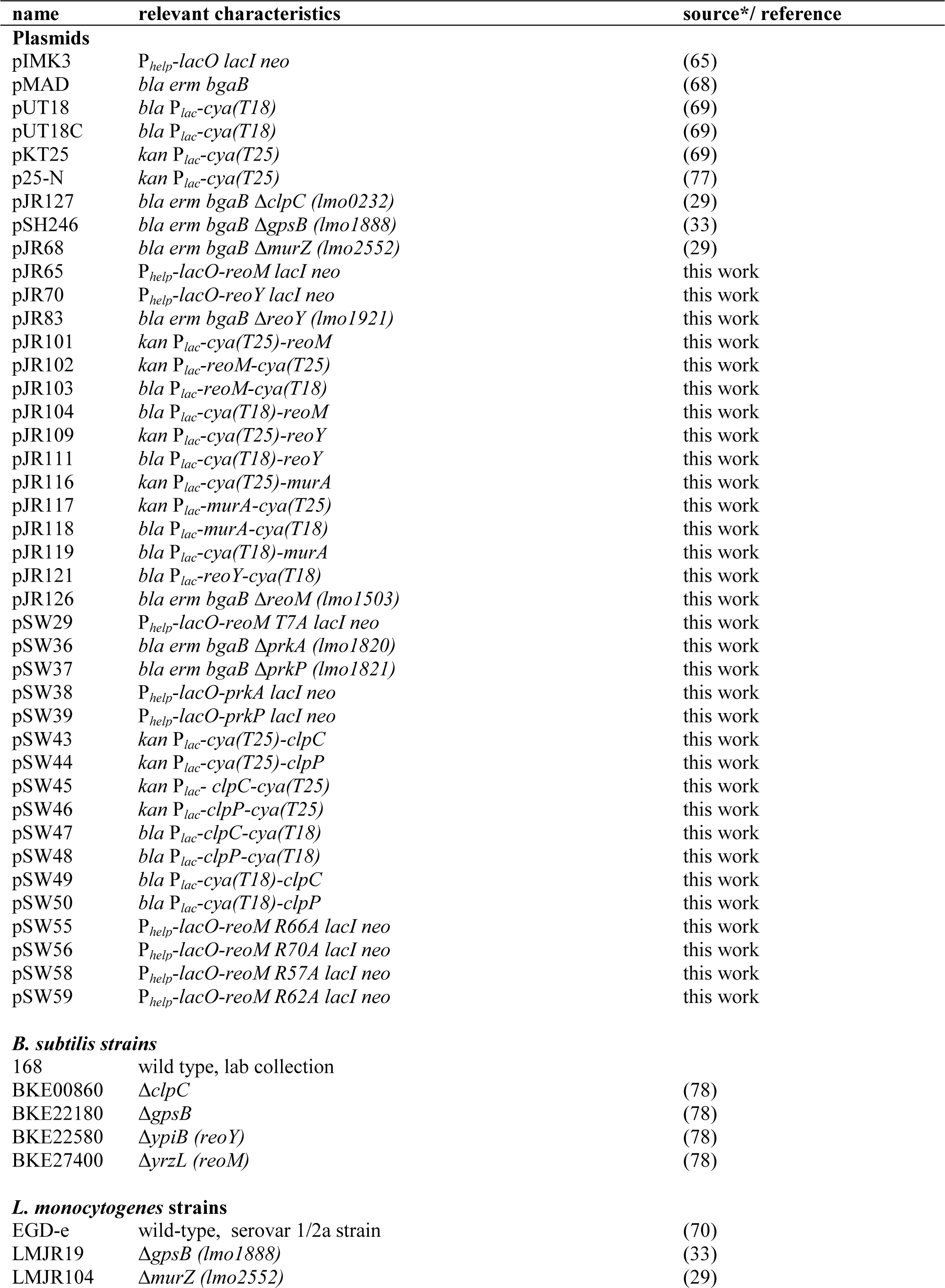
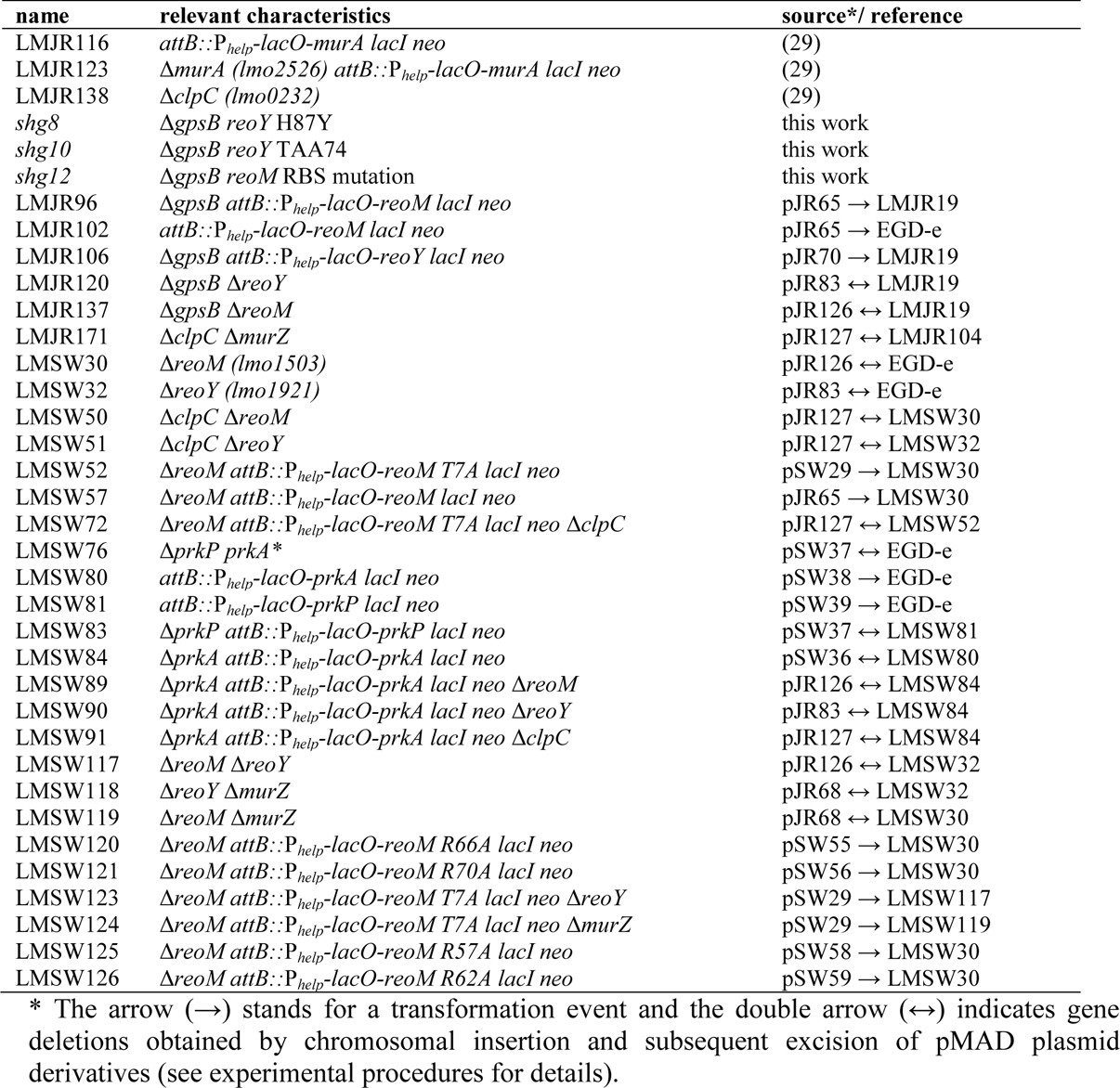
Plasmids and strains used in this study

| name | relevant characteristics | source*/ reference |
| --- | --- | --- |
| **Plasmids** |  |  |
| pIMK3 | *P<sub>help</sub>-lacO lacI neo* | (65) |
| pMAD | *bla erm bgaB* | (68) |
| pUT18 | *bla P<sub>lac</sub>-cya(T18)* | (69) |
| pUT18C | *bla P<sub>lac</sub>-cya(T18)* | (69) |
| pKT25 | *kan P<sub>lac</sub>-cya(T25)* | (69) |
| p25-N | *kan P<sub>lac</sub>-cya(T25)* | (77) |
| pJR127 | *bla erm bgaB ΔclpC (lmo0232)* | (29) |
| pSH246 | *bla erm bgaB ΔgpsB (lmo1888)* | (33) |
| pJR68 | *bla erm bgaB ΔmurZ (lmo2552)* | (29) |
| pJR65 | *P<sub>help</sub>-lacO-reoM lacI neo* | this work |
| pJR70 | *P<sub>help</sub>-lacO-reoY lacI neo* | this work |
| pJR83 | *bla erm bgaB ΔreoY (lmo1921)* | this work |
| pJR101 | *kan P<sub>lac</sub>-cya(T25)-reoM* | this work |
| pJR102 | *kan P<sub>lac</sub>-reoM-cya(T25)* | this work |
| pJR103 | *bla P<sub>lac</sub>-reoM-cya(T18)* | this work |
| pJR104 | *bla P<sub>lac</sub>-cya(T18)-reoM* | this work |
| pJR109 | *kan P<sub>lac</sub>-cya(T25)-reoY* | this work |
| pJR111 | *bla P<sub>lac</sub>-cya(T18)-reoY* | this work |
| pJR116 | *kan P<sub>lac</sub>-cya(T25)-murA* | this work |
| pJR117 | *kan P<sub>lac</sub>-murA-cya(T25)* | this work |
| pJR118 | *bla P<sub>lac</sub>-murA-cya(T18)* | this work |
| pJR119 | *bla P<sub>lac</sub>-cya(T18)-murA* | this work |
| pJR121 | *bla P<sub>lac</sub>-reoY-cya(T18)* | this work |
| pJR126 | *bla erm bgaB ΔreoM (lmo1503)* | this work |
| pSW29 | *P<sub>help</sub>-lacO-reoM T7A lacI neo* | this work |
| pSW36 | *bla erm bgaB ΔprkA (lmo1820)* | this work |
| pSW37 | *bla erm bgaB ΔprkP (lmo1821)* | this work |
| pSW38 | *P<sub>help</sub>-lacO-prkA lacI neo* | this work |
| pSW39 | *P<sub>help</sub>-lacO-prkP lacI neo* | this work |
| pSW43 | *kan P<sub>lac</sub>-cya(T25)-clpC* | this work |
| pSW44 | *kan P<sub>lac</sub>-cya(T25)-clpP* | this work |
| pSW45 | *kan P<sub>lac</sub>- clpC-cya(T25)* | this work |
| pSW46 | *kan P<sub>lac</sub>-clpP-cya(T25)* | this work |
| pSW47 | *bla P<sub>lac</sub>-clpC-cya(T18)* | this work |
| pSW48 | *bla P<sub>lac</sub>-clpP-cya(T18)* | this work |
| pSW49 | *bla P<sub>lac</sub>-cya(T18)-clpC* | this work |
| pSW50 | *bla P<sub>lac</sub>-cya(T18)-clpP* | this work |
| pSW55 | *P<sub>help</sub>-lacO-reoM R66A lacI neo* | this work |
| pSW56 | *P<sub>help</sub>-lacO-reoM R70A lacI neo* | this work |
| pSW58 | *P<sub>help</sub>-lacO-reoM R57A lacI neo* | this work |
| pSW59 | *P<sub>help</sub>-lacO-reoM R62A lacI neo* | this work |
| ***B. subtilis* strains** |  |  |
| 168 | wild type, lab collection |  |
| BKE00860 | *ΔclpC* | (78) |
| BKE22180 | *ΔgpsB* | (78) |
| BKE22580 | *ΔypiB (reoY)* | (78) |
| BKE27400 | *ΔyrzL (reoM)* | (78) |
| ***L. monocytogenes* strains** |  |  |
| EGD-e | wild-type, serovar 1/2a strain | (70) |
| LMJR19 | *ΔgpsB (lmo1888)* | (33) |
| LMJR104 | *ΔmurZ (lmo2552)* | (29) |
| LMJR116 | *attB::P<sub>help</sub>-lacO-murA lacI neo* | (29) |
| LMJR123 | *ΔmurA (lmo2526) attB::P<sub>help</sub>-lacO-murA lacI neo* | (29) |
| LMJR138 | *ΔclpC (lmo0232)* | (29) |
| *shg8* | *ΔgpsB reoY H87Y* | this work |
| *shg10* | *ΔgpsB reoY TAA74* | this work |
| *shg12* | *ΔgpsB reoM* RBS mutation | this work |
| LMJR96 | *ΔgpsB attB::P<sub>help</sub>-lacO-reoM lacI neo* | pJR65 → LMJR19 |
| LMJR102 | *attB::P<sub>help</sub>-lacO-reoM lacI neo* | pJR65 → EGD-e |
| LMJR106 | *ΔgpsB attB::P<sub>help</sub>-lacO-reoY lacI neo* | pJR70 → LMJR19 |
| LMJR120 | *ΔgpsB ΔreoY* | pJR83 ↔ LMJR19 |
| LMJR137 | *ΔgpsB ΔreoM* | pJR126 ↔ LMJR19 |
| LMJR171 | *ΔclpC ΔmurZ* | pJR127 ↔ LMJR104 |
| LMSW30 | *ΔreoM (lmo1503)* | pJR126 ↔ EGD-e |
| LMSW32 | *ΔreoY (lmo1921)* | pJR83 ↔ EGD-e |
| LMSW50 | *ΔclpC ΔreoM* | pJR127 ↔ LMSW30 |
| LMSW51 | *ΔclpC ΔreoY* | pJR127 ↔ LMSW32 |
| LMSW52 | *ΔreoM attB::P<sub>help</sub>-lacO-reoM T7A lacI neo* | pSW29 → LMSW30 |
| LMSW57 | *ΔreoM attB::P<sub>help</sub>-lacO-reoM lacI neo* | pJR65 → LMSW30 |
| LMSW72 | *ΔreoM attB::P<sub>help</sub>-lacO-reoM T7A lacI neo ΔclpC* | pJR127 ↔ LMSW52 |
| LMSW76 | *ΔprkP prkA\** | pSW37 ↔ EGD-e |
| LMSW80 | *attB::P<sub>help</sub>-lacO-prkA lacI neo* | pSW38 → EGD-e |
| LMSW81 | *attB::P<sub>help</sub>-lacO-prkP lacI neo* | pSW39 → EGD-e |
| LMSW83 | *ΔprkP attB::P<sub>help</sub>-lacO-prkP lacI neo* | pSW37 ↔ LMSW81 |
| LMSW84 | *ΔprkA attB::P<sub>help</sub>-lacO-prkA lacI neo* | pSW36 ↔ LMSW80 |
| LMSW89 | *ΔprkA attB::P<sub>help</sub>-lacO-prkA lacI neo ΔreoM* | pJR126 ↔ LMSW84 |
| LMSW90 | *ΔprkA attB::P<sub>help</sub>-lacO-prkA lacI neo ΔreoY* | pJR83 ↔ LMSW84 |
| LMSW91 | *ΔprkA attB::P<sub>help</sub>-lacO-prkA lacI neo ΔclpC* | pJR127 ↔ LMSW84 |
| LMSW117 | *ΔreoM ΔreoY* | pJR126 ↔ LMSW32 |
| LMSW118 | *ΔreoY ΔmurZ* | pJR68 ↔ LMSW32 |
| LMSW119 | *ΔreoM ΔmurZ* | pJR68 ↔ LMSW30 |
| LMSW120 | *ΔreoM attB::P<sub>help</sub>-lacO-reoM R66A lacI neo* | pSW55 → LMSW30 |
| LMSW121 | *ΔreoM attB::P<sub>help</sub>-lacO-reoM R70A lacI neo* | pSW56 → LMSW30 |
| LMSW123 | *ΔreoM attB::P<sub>help</sub>-lacO-reoM T7A lacI neo ΔreoY* | pSW29 → LMSW117 |
| LMSW124 | *ΔreoM attB::P<sub>help</sub>-lacO-reoM T7A lacI neo ΔmurZ* | pSW29 → LMSW119 |
| LMSW125 | *ΔreoM attB::P<sub>help</sub>-lacO-reoM R57A lacI neo* | pSW58 → LMSW30 |
| LMSW126 | *ΔreoM attB::P<sub>help</sub>-lacO-reoM R62A lacI neo* | pSW59 → LMSW30 |

\* The arrow (→) stands for a transformation event and the double arrow (↔) indicates gene deletions obtained by chromosomal insertion and subsequent excision of pMAD plasmid derivatives (see experimental procedures for details).

### General methods, manipulation of DNA and oligonucleotide primers

Standard methods were used for transformation of *E. coli* and isolation of plasmid DNA (63). Transformation of *L. monocytogenes* was carried out as described by others (65). Restriction and ligation of DNA was performed according to the manufactureŕs instructions. All primer sequences are listed in Tab. 4.

**Table 4:**
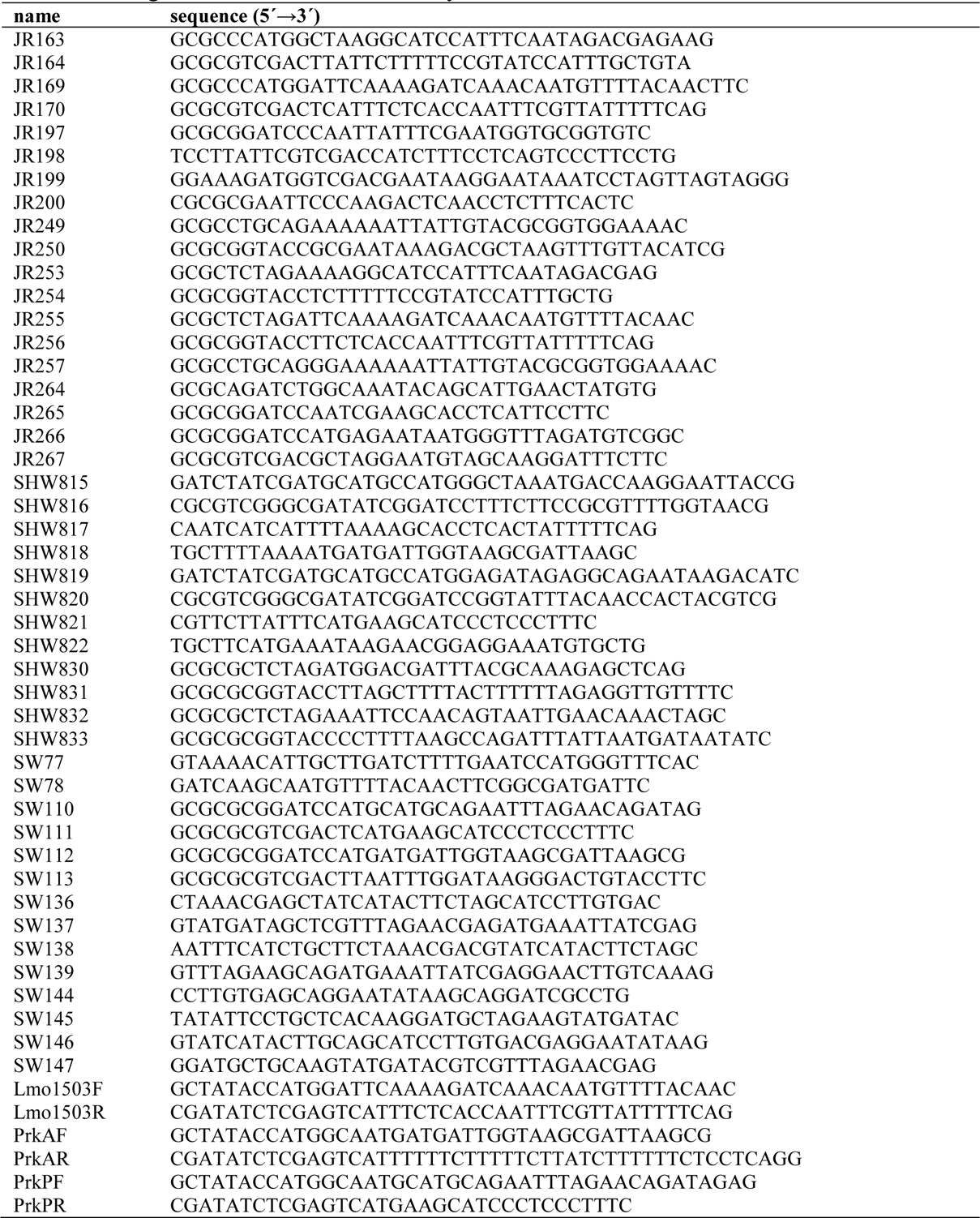
Oligonucleotides used in this study.

### Construction of plasmids for recombinant protein expression

The plasmids for expressing recombinant versions of ReoM, PrkA-KD and PrkP were prepared by first amplifying the corresponding genes (*reoM*, *lmo1820* and *lmo1821*) from *L. monocytogenes* EGD-e genomic DNA using primer pairs Lmo1503F/Lmo1503R, PrkAF/PrkAR, and PrkPF/PrkPR, respectively. The PCR products were individually ligated between the *Nco*I and *Xho*I sites of pETM11 (66). All mutagenesis was carried out using the Quikchange protocol and the correct sequence of each plasmid and mutant constructed was verified by Sanger DNA sequencing (Eurofins Genomics).

### Construction of plasmids for generation of *L. monocytogenes* strains

Plasmid pJR65 was constructed for the inducible expression of *reoM*. To this end, the *reoM* open reading frame was amplified by PCR using the oligonucleotides JR169/JR170 and cloned into pIMK3 using NcoI/SalI. The T7A mutation were introduced into *reoM* of plasmid pJR65 by quickchange mutagenesis using the primer pair SW77/SW78, yielding pSW29. The R57A, R62A R66A and R70A, mutations were introduced into pJR65 in the same way, but using primer pairs SW144/SW145, SW146/SW147, SW136/SW137 and SW138/SW139, respectively.

Plasmid pJR70 was constructed for inducible *reoY* expression. For this purpose, *reoY* was amplified using the primer pair JR163/JR164 and cloned into pIMK3 using NcoI/SalI.

Plasmid pSW38, for IPTG-inducible *prkA* expression, was constructed by amplification of *prkA* using the oligonucleotides SW112/SW113 and the subsequent cloning of the generated fragment into pIMK3 using BamHI/SalI. Plasmid pSW39, for IPTG-controlled expression of *prkP*, was constructed analogously, but using the oligonucleotides SW110/SW111 as the primers.

For construction of plasmid pJR83, facilitating deletion of *reoY*, fragments encompassing ∼800 bp up- and down-stream of *reoY* were amplified by PCR with the primer pairs JR197/JR198 and JR199/JR200. Both fragments were spliced together by splicing by overlapping extension (SOE) PCR and cloned into pMAD using BamHI/EcoRI.

Plasmid pJR126 was generated for deletion of *reoM*. Fragments up- and down-stream of *reoM* were PCR amplified using the primers JR264/JR265 and JR266/JR267, repsectively. Both fragments were cut with BamHI, fused together by ligation and the desired fragment was amplified from the ligation mixture by PCR using the primers JR264/JR267 and then cloned into pMAD using BglII/SalI.

Plasmid pSW36 was constructed to delete the *prkA* gene. Fragments up- and down-stream of *prkA* were amplified in separate PCRs using the primer pairs SHW819/SHW821 and SHW820/SHW822, respectively. Both fragments were fused together by SOE-PCR and inserted into pMAD by restriction free cloning (67). Plasmid pSW37, facilitating deletion of *prkP*, was constructed in a similar manner. Up- and down-stream fragments of *prkP* were amplified using the primer pairs SHW815/SHW817 and SHW816/SHW818 and fused together by SOE-PCR. The resulting fragment was inserted into pMAD by restriction free cloning.

Derivatives of pIMK3 were introduced into *L. monocytogenes* strains by electroporation and clones were selected on BHI agar plates containing kanamycin. Plasmid insertion at the *attB* site of the tRNA^Arg^ locus was verified by PCR. Plasmid derivatives of pMAD were transformed into the respective *L. monocytogenes* recipient strains and genes were deleted as described elsewhere (68). All gene deletions were confirmed by PCR.

### Construction of bacterial two hybrid plasmids

The *reoM* (JR255/JR256), *reoY* (JR253/JR254), *clpC* (SHW830/831) and *clpP* (SHW832/833) genes were amplified using the primer pairs given in brackets and cloned into pUT18, pUT18C, pKT25 and p25-N plasmids using XbaI/KpnI. The *murA* gene was amplified using the oligonucleotides JR249/JR250 for cloning into pKT25 and p25-N using PstI/KpnI or using the JR257/JR250 primer pair for cloning into pUT18 and pUT18C using the same restriction enzymes.

### Bacterial two hybrid experiments

Plasmids carrying genes fused to T18- or the T25-fragments of the *Bordetella pertussis* adenylate cyclase were co-transformed into *E. coli* BTH101 (69) and transformants were selected on LB agar plates containing ampicillin (100 µg ml^-1^), kanamycin (50 µg ml^-1^), X-Gal (0.004%) and IPTG (0.1 mM). Agar plates were photographed after 48 h of incubation at 30°C.

### Genome sequencing

A total of 1 ng of genomic DNA was used for library generation by the Nextera XT DNA Library Prep Kit according to the manufacturer’s recommendations (Illumina). Sequencing was carried out on a MiSeq benchtop sequencer and performed in paired-end modes (2 x 300 bp) using a MiSeq Reagent Kit v3 cartridge (600-cycle kit). Sequencing reads were mapped to the reference genome *L. monocytogenes* EGD-e (NC_003210.1) (70) by utilizing the Geneious software (Biomatters Ltd.). Variants, representing putative suppressor mutations, were identified using the Geneious SNP finder tool. Genome sequences of *shg8*, *shg10*, *shg12* and LMSW76 were deposited at ENA under study number PRJEB35110 and sample accession numbers ERS3927571 (SAMEA6127277), ERS3927572 (SAMEA6127278), ERS3927573 (SAMEA6127279), and ERS3967687 (SAMEA6167687) respectively.

### Isolation of cellular proteins and Western blotting

Cells were harvested by centrifugation (13,000 rpm, 1 min in a table-top centrifuge), washed with ZAP buffer (10 mM Tris.HCl pH7.5, 200 mM NaCl), resuspended in 1 ml ZAP buffer also containing 1 mM PMSF and disrupted by sonication. Centrifugation was used to remove cellular debris and the supernatant was used as total cellular protein extract. Sample aliquots were separated by standard SDS polyacrylamide gel electrophoresis. Gels were transferred onto positively charged polyvinylidene fluoride membranes by semi-dry transfer. ClpC, DivIVA, GlmS, IlvB and MurA were immune-stained using a polyclonal rabbit antiserum recognizing *B. subtilis* ClpC (29), DivIVA (71), GlmS, IlvB (40) and MurAA (27) as the primary antibody and an anti-rabbit immunoglobulin G conjugated to horseradish peroxidase as the secondary one. The ECL chemiluminescence detection system (Thermo Scientific) was used for detection of the peroxidase conjugates on the PVDF membrane in a chemiluminescence imager (Vilber Lourmat). For depletion of PrkA, PrkA depletion strains were grown overnight in the presence of 1 mM IPTG and then again inoculated in BHI broth containing 1 mM IPTG to an OD_600_=0.05 and grown for 3 h at 37°C. Subsequently, cells were centrifuged, washed and reinoculated in BHI broth without IPTG at the same OD_600_ as before centrifugation. Finally, cells were harvested after 3.5 more hours of growth at 37°C and cellular proteins were isolated.

### Microscopy

Cytoplasmic membranes of exponentially growing bacteria were stained through addition of 1 µl of nile red solution (100 µg ml^-1^ in DMSO) to 100 µl of culture. Images were taken with a Nikon Eclipse Ti microscope coupled to a Nikon DS-MBWc CCD camera and processed using the NIS elements AR software package (Nikon) or ImageJ. Scanning electron microscopy was performed essentially as described earlier (64).

### Recombinant protein purification

All proteins were expressed in *E. coli* BL21 (DE3) cells. Cell cultures were grown at 37°C in LB liquid media supplemented with 50 µg mL^-1^ kanamycin to an OD_600_ 0.6-0.8 before expression was induced by the addition of IPTG to a final concentration of 0.4 mM IPTG. The cultures were incubated at 20°C overnight before the cells from 2 L of cell culture were harvested by centrifugation at 3500 x g for 30 minutes. The cell pellets were resuspended in 70 mL of buffer A (50 mM Tris.HCl, pH 8, 300 mM NaCl, 10 mM imidazole) with 500 Kunitz units of DNase I and 1 mL Roche complete protease inhibitor cocktail at 25x working concentration. The cells were lysed by sonication, centrifuged at 19000 x g for 20 minutes and the supernatant was filtered using a 0.45 µm filter. The clarified cell lysate was loaded onto a 5 mL Ni-NTA superflow cartridge (Qiagen), washed with buffer A, and bound proteins were eluted with 50 mM Tris.HCl, pH 8, 300 mM NaCl, 250 mM imidazole. The His_6_-tag of PrkA-KD was cleaved with His-tagged TEV protease (1 mg TEV for 20 mg of protein) at 4 °C during an overnight dialysis against a buffer of 50 mM Tris.HCl, pH 8, 300 mM NaCl, 10 mM imidazole, 1 mM DTT; TEV cleavage of ReoM was conducted as above except the dialysis was carried out at 20 °C. The proteolysis reaction products were then passed over a 5 mL Ni-NTA superflow cartridge (Qiagen) to remove TEV and uncleaved protein. The proteins that did not bind to the Ni-NTA column were concentrated and loaded onto either a Superdex 75 XK16/60 (GE Healthcare) (ReoM) or a Superdex 200 XK16/60 (GE Healthcare) (PrkA-KD and PrkP) equilibrated with 10 mM Na-HEPES, pH 8, 100 mM NaCl for size exclusion chromatography. Fractions from the gel filtration were analysed for purity by SDS-PAGE, concentrated to 20-40 mg mL^-1^, and small aliquots were snap-frozen in liquid nitrogen for storage at −80°C.

### X-ray crystallography and ReoM structure determination

For ReoM, 23 mg mL^-1^ of protein in 10 mM Na-HEPES pH 8, 100 mM NaCl was subjected to crystallisation by sparse matrix screening using a panel of commercial crystallisation screens. 100 and 200 nL drops of protein and 100 nL of screen solution were dispensed into 96 well MRC crystallization plates (Molecular Dimensions) by a Mosquito (TTP Labtech) liquid handling robot and the crystallisation plates were stored at a constant temperature of 20°C. The crystals that grew and were subsequently used for diffraction experiments were formed in 0.1 M phosphate/citrate pH 4.2, 0.2 M lithium sulfate, 20 % w/v PEG 1000 from the JCSG + screen and were mounted onto rayon loops directly from the crystallization drops and cryo-cooled in liquid nitrogen.

Diffraction data were collected on beamline I03 at the Diamond Light Source (DLS) synchrotron. Diffraction images were integrated in MOSFLM (72) and scaled and merged with AIMLESS (73). The initial model was generated by molecular replacement in PHASER (74) using the dimeric, 20-conformer ensemble model (PDBid 5US5) of IreB solved by nuclear magnetic resonance (45) as a search model. The final model was produced by iterative cycles of model building in COOT (75) with refinement in REFMAC (76) until convergence. The diffraction data collection and model refinement statistics are summarised in Tab. 2.

### Protein phosphorylation and dephosphorylation

The effect of phosphorylation and dephosphorylation on ReoM and PrkA-KD proteins was analysed by 20% non-denaturing PAGE. Phosphorylation reactions consisted of 18.5 µM ReoM, 3.7 µM PrkA-KD, 5 mM ATP and 5 mM MgCl_2_, diluted in 10 mM HEPES.HCl pH 8.0 and 100 mM NaCl. Dephosphorylation reactions consisted of 37 µM P-ReoM, 3.7 µM PrkA-KD, 18.5 µM PrkP and 1 mM MnCl_2_, diluted in 10 mM HEPES.HCl pH 8.0 and 100 mM NaCl. In each case controls were loaded at the same concentrations. The reactions were incubated at 37 °C for 20 minutes prior to electrophoresis at 200 V for 2.5 hours on ice.

### Isolation of phosphorylated ReoM

Phosphorylation reactions consisted of 37 µM ReoM, 3.7 µM PrkA-KD, 5 mM ATP and 5 mM MgCl_2_, diluted in 10 mM HEPES.HCl pH 8.0 and 100 mM NaCl, to a total volume of 5 mL. The protein mix was loaded onto a PD 10 desalting column to remove excess ATP and protein fractions were loaded onto a MonoQ 5/50 GL column. Buffer A consisted of 10 mM HEPES.HCl pH 8.0 and 100 mM NaCl and buffer B was 10 mM HEPES.HCl pH 8.0 and 1M NaCl. Bound proteins were eluted over 25 mL with a 15-35% gradient of buffer B.

### Liquid Chromatography-Mass Spectrometry

All liquid chromatography-mass spectrometry (LC-MS) analyses were performed using an Agilent 6530 Q-TOF instrument with electrospray ionisation (ESI) in positive ion mode, coupled to an Agilent 1260 Infinity II LC system, utilizing mobile phase of 0.1% (v/v) formic acid in LC-MS grade water (A) and acetonitrile (B). Prior to peptide mapping, 10 μL of purified proteins (∼1 mg/ml) were digested using Smart Digest Soluble Trypsin Kit (Thermo Fisher Scientific) according to the manufacturer’s guidelines. Tryptic peptides and intact protein samples were extracted using HyperSep Spin Tip SPE C18 and C8 tips, respectively (ThermoFisher Scientific) before analysis. For phosphosite analysis, 10 μL of digest was injected onto an Acclaim RSLC 120 C18 column (Thermo Fisher Scientific, 2.1 x 100mm, 2.2 µm, 120 Å) for reversed phase separation at 60°C and 0.4 ml/min, over a linear gradient of 5-40% B over 25 min, 40-90% B over 8 min followed by equilibration at 5% B for 7 min. ESI source conditions were nebuliser pressure of 45 psig, drying gas flow of 5 L/min and gas temperature of 325°C. Sheath gas temperature of 275°C and gas flow of 12 L/min, capillary voltage of 4000V and nozzle voltage of 300V were also applied. Mass spectra were acquired using MassHunter Acquisition software (version B.08.00) over the 100-3000 m/z range, at a rate of 5 spectra/s and 200 ms/spectrum, using standard mass range mode (3200 m/z) with extended dynamic range (2 GHz) and collection of both centroid and profile data. MS/MS fragmentation spectra were acquired over the 100-3000 m/z range, at a rate of 3 spectra/s and 333.3 ms/spectrum. For intact protein analysis,10 μL of desalted protein (∼1 mg/ml) was injected onto a Zorbax 300Å Stable Bond C8 column (Agilent Technologies, 4.6 x 50 mm, 3.5 μM) for reversed phase separation at 60°C and 0.4 mL/min, over a linear gradient of 15-75% B over 14 min, 75-100% B over 11 min followed by post-run equilibration at 15% B for 10 min. ESI source conditions were nebuliser pressure of 45 psig, drying gas flow of 5 L/min and source gas temperature of 325°C were applied. Sheath gas temperature of 400°C and gas flow of 11 L/min, capillary voltage of 3500V and nozzle voltage of 2000V were also used. Mass spectra were acquired using MassHunter Acquisition software (version B.08.00) over a mass range of 100-3000 m/z, at a rate of 1 spectra/s and 1000 ms/spectrum in extended mass range (20000 m/z) at 1 GHz. Acquired MS and MS/MS spectra were analysed using Agilent MassHunter BioConfirm software (version B.10.00) for identification of phosphorylated residues and subsequent intact mass determination with processing of raw data using maximum entropy deconvolution.

### Analytical size exclusion chromatography

Purified ReoM and P-ReoM proteins were run individually on a Superdex 200 Increase 10/300 GL column. 100 µl samples at 1.5 mg/mL were injected onto a column equilibrated in 10 mM HEPES.HCl pH 8.0 and 100 mM NaCl, with a flow of 0.75 mL/min.

## ACKNOWLEDGEMENTS

This work was funded by DFG grants HA 6830/1-1 and HA 6830/1-2 and a grant of the Fonds der Chemischen Industrie to SH. ZR is funded by a UK BBSRC DTP studentship to RJL (BB/M011186/1). We acknowledge Diamond Light Source for time on beamline I03 under proposal MX-18598 and Dr. Arnaud Basle for help with X-ray data collection. We thank Ulrich Nübel (Braunschweig) and Janina Döhling for help with some experiments and Petra Kaiser for technical assistance. We would like to thank the National BioResource Project (NIG, Japan): *B. subtilis* for sharing *B. subtilis* mutant strains. The co-ordinates and structure factors for the crystal structure of ReoM have been deposited at PDBe with accession code 6TIF.

## AUTHOR CONTRIBUTIONS

SW, ZJR, JR, CEJ, LM, RJL and SH designed the experiments. SW, ZJR, JR, CEJ and LM performed the experimental work. SW, ZJR, JR, CEJ, LM, RJL and SH interpreted the data. RJL and SH wrote the manuscript.

