## Supplementary material for "PrkA controls peptidoglycan biosynthesis through the essential phosphorylation of ReoM": Suplementary Figures S1-S13

by

**Sabrina Wamp, Zoe J.Rutter, Jeanine Rismondo, Claire E. Jennings, Lars Möller, Richard J. Lewis and Sven Halbedel**

**Figure S1:** Overexpression of *reoM* but not *reoY* affects growth of the *L. monocytogenes* Δ*gpsB* mutant.

**Figure S2:** Effect of *reoM* and *reoY* deletions on cell morphology.

**Figure S3:** ReoM, ReoY and MurZ operate in the same pathway.

**Figure S4:** DivIVA stability in *L. monocytogenes* Δ*clpC*, Δ*reoM* and Δ*reoY* mutants.

**Figure S5:** Effect of *reoM* and *reoY* deletions on accumulation of other ClpC substrates in *B. subtilis*.

**Figure S6**: LC-MS analysis of intact ReoM.

**Figure S7:** LC-MS analysis of ReoM T7A.

**Figure S8**: Dephosphorylation of P-ReoM by PrkP.

**Figure S9**: Dephosphorylation of P-PrkA-KD by PrkP.

**Figure S10:** ReoM and P-ReoM have the same oligomeric state.

**Figure S11**: Lethality of ReoM R57A and R62A subtitutions.

**Figure S12**: A possible conformational change of the flexible ReoM N-terminus induced by phosphorylation.

**Figure S13:** Bacterial two hybrid experiment showing interactions between MurA, ReoM, ReoY, ClpC and ClpP.

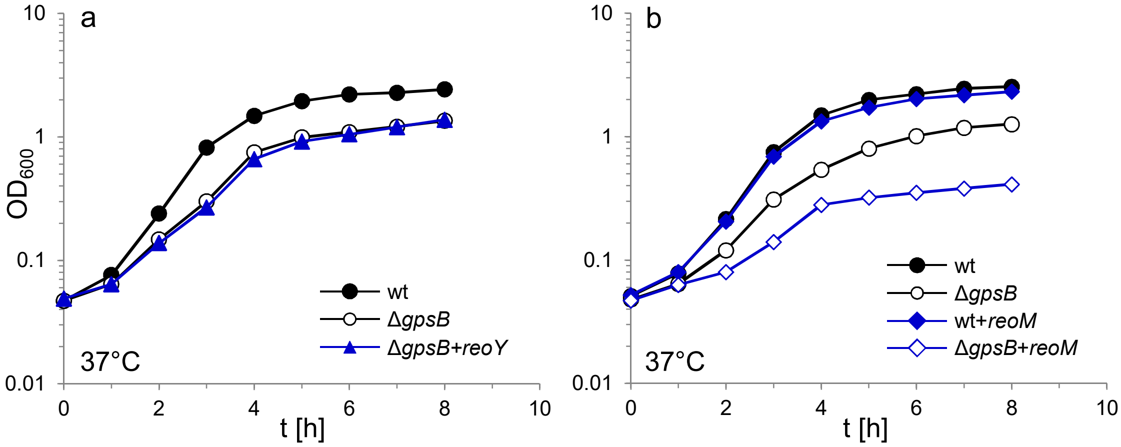


**Figure S1:** Overexpression of *reoM* but not *reoY* affects growth of the *L. monocytogenes* Δ*gpsB* mutant.
(A) Growth of *L. monocytogenes* strains EGD-e (wt), LMJR19 (Δ*gpsB*) and LMJR106 (Δ*gpsB*+*reoY*) in BHI broth containing 1 mM IPTG at 37°C. (B) Growth of strains EGD-e (wt), LMJR19 (Δ*gpsB*), LMJR102 (wt+*reoM*) and LMJR96 (Δ*gpsB*+*reoM*) in BHI broth containing 1 mM IPTG at 37°C. The experiments were performed twice and one representative result is shown.

Wamp *et al.*, Fig. S1

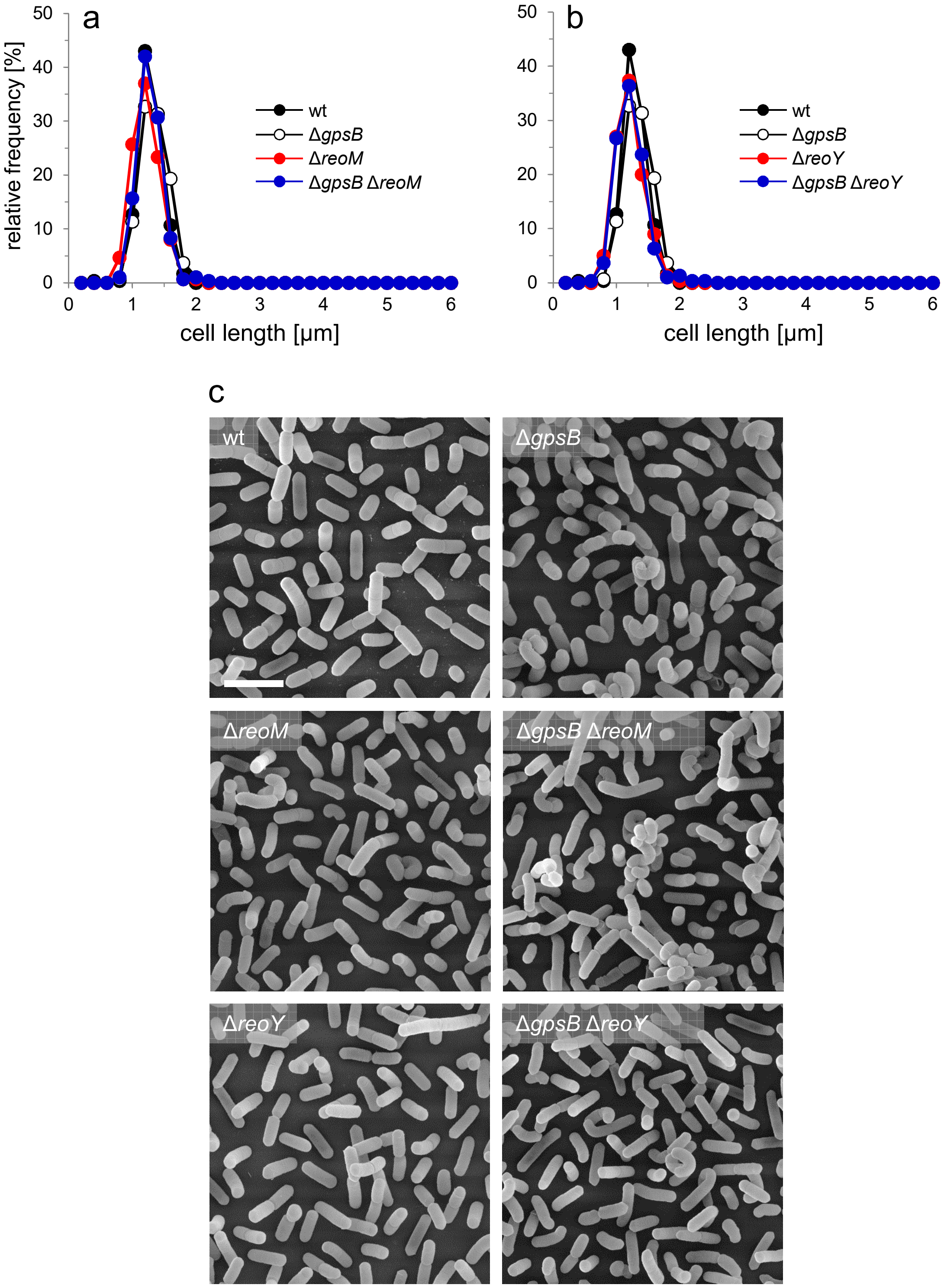


**Figure S2:** Effect of *reoM* and *reoY* deletions on cell morphology.
Strains EGD-e (wt), LMJR19 (Δ*gpsB*), LMSW30 (Δ*reoM*), LMSW32 (Δ*reoY*), LMJR137 (Δ*gpsB* Δ*reoM*), and LMJR120 (Δ*gpsB* Δ*reoY*) were grown in BHI broth at 37°C to mid-logarithmic growth phase and cell lengths were measured after nile red staining and fluorescence microscopy. Diagrams show the effect of *reoM* (A) and *reoY* (B) deletions on cell lengths of wild type and Δ*gpsB* mutant cells as cell length frequency distributions determined on 300 cells per strain from a single culture. Results of one out of three experiments are shown. (C) Scanning electron microscopy of the same strains as in panel A and B. Strains were grown to mid-logarithmic growth phase in BHI at 37°C and subjected to chemical fixation and subsequent electron microscopy as described in the experimental procedures section. Scale bar: 2 µm.

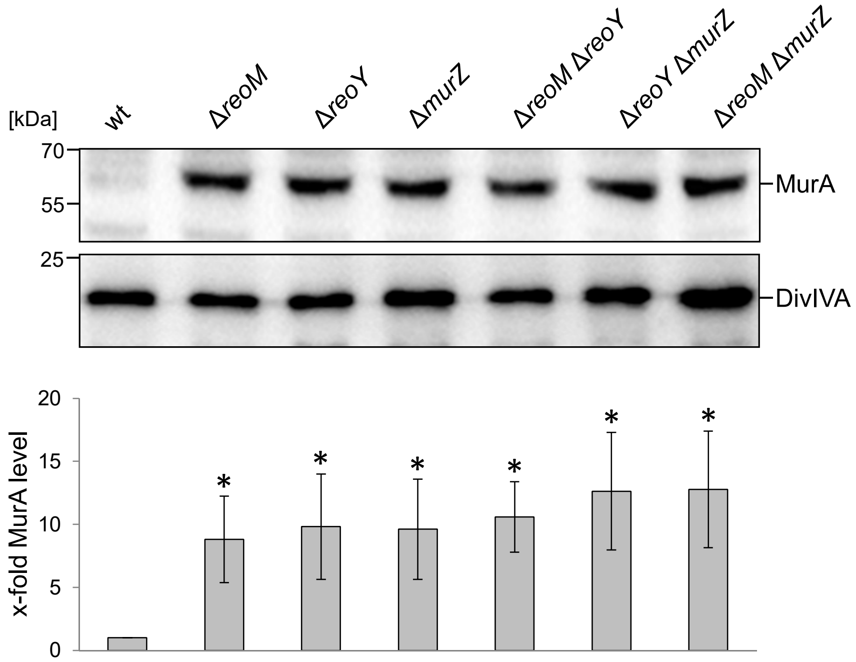


**Figure S3:** ReoM, ReoY and MurZ operate in the same pathway.
Western blot showing MurA amounts in *L. monocytogenes* strains EGD-e (wt), LMSW30 (Δ*reoM*), LMSW32 (Δ*reoY*), LMJR104 (Δ*murZ*), LMSW117 (Δ*reoM* Δ*reoY),* LMSW118 (Δ*reoY* Δ*murZ)* and LMSW119 (Δ*reoM* Δ*murZ*, top*)*. A parallel Western blot using an α-DivIVA antiserum was used as loading control (middle panel). Strains were grown to mid-logarithmic growth phase for isolation of cellular proteins. Quantification of MurA signals by densitometry (bottom). Average values and standard deviations calculated from three independent experiments are shown. Asterisks indicate statistically significant differences compared to wild type ($P<0.05$, *t*-test).

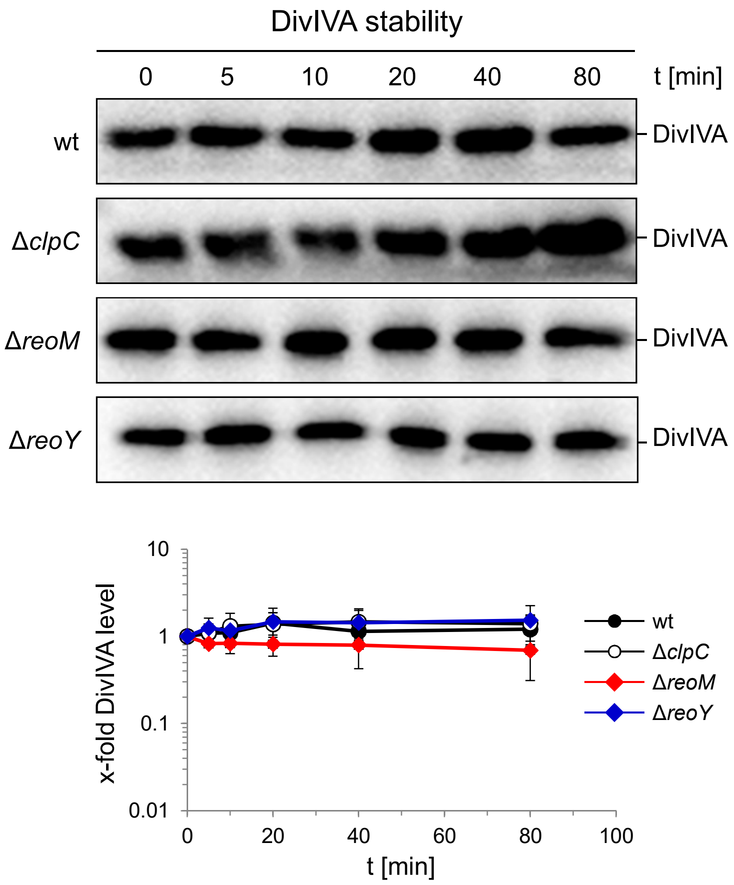


**Figure S4:** DivIVA stability in *L. monocytogenes* Δ*clpC*, Δ*reoM* and Δ*reoY* mutants.
Western blots following DivIVA levels after chloramphenicol treatment. *L. monocytogenes* strains EGD-e (wt), LMJR138 (Δ*clpC*), LMSW30 (Δ*reoM*) and LMSW32 (Δ*reoY*) were grown to an $OD_{600}$ of 1.0 and 100 µg/ml chloramphenicol was added to block protein biosynthesis. Samples were taken before chloramphenicol addition and after several time intervals to analyze DivIVA levels. DivIVA signals were quantified by densitometry and average values and standard deviations are shown (n=3).

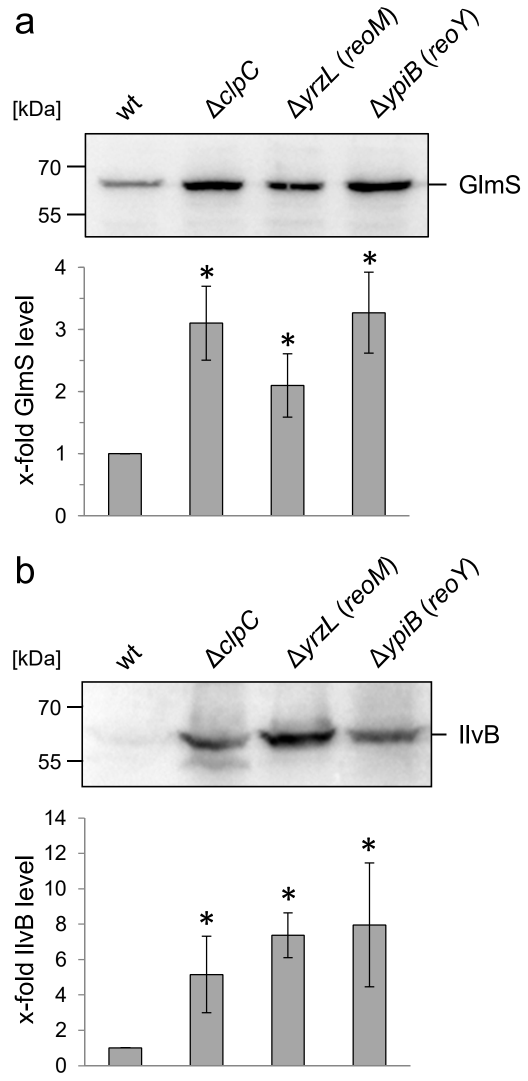


**Figure S5:** Effect of *reoM* and *reoY* deletions on accumulation of other ClpC substrates in *B. subtilis*.
(A) Western blot showing GlmS amounts in *B. subtilis* strains 168 (wt), BKE00860 (Δ*clpC*), BKE27400 (Δ*yrzL/reoM*) and BKE22580 (Δ*ypiB/reoY*, top). Strains were grown to mid-logarithmic growth phase for isolation of cellular proteins. Quantification of GlmS signals by densitometry (bottom). (B) Western blot showing IlvB amounts in the same set of strains as in panel A (top). Quantification of IlvB signals by densitometry (bottom). Average values and standard deviations calculated from three independent experiments are shown. Asterisks indicate statistically significant differences ($P$<0.05, *t*-test). A loading control is shown in Fig. 2C.

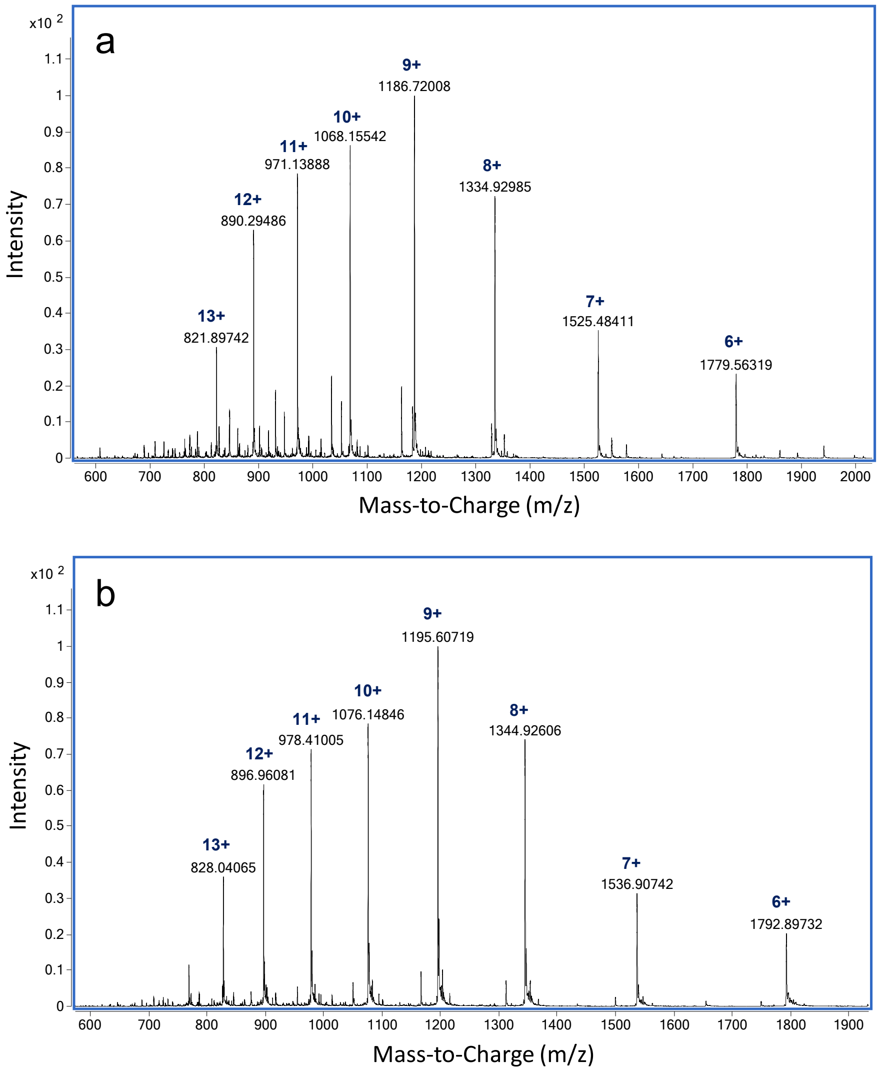


**Figure S6**: LC-MS analysis of intact ReoM.
ESI-QTOF raw data for A) unmodified ReoM and B) mono-phosphorylated ReoM.

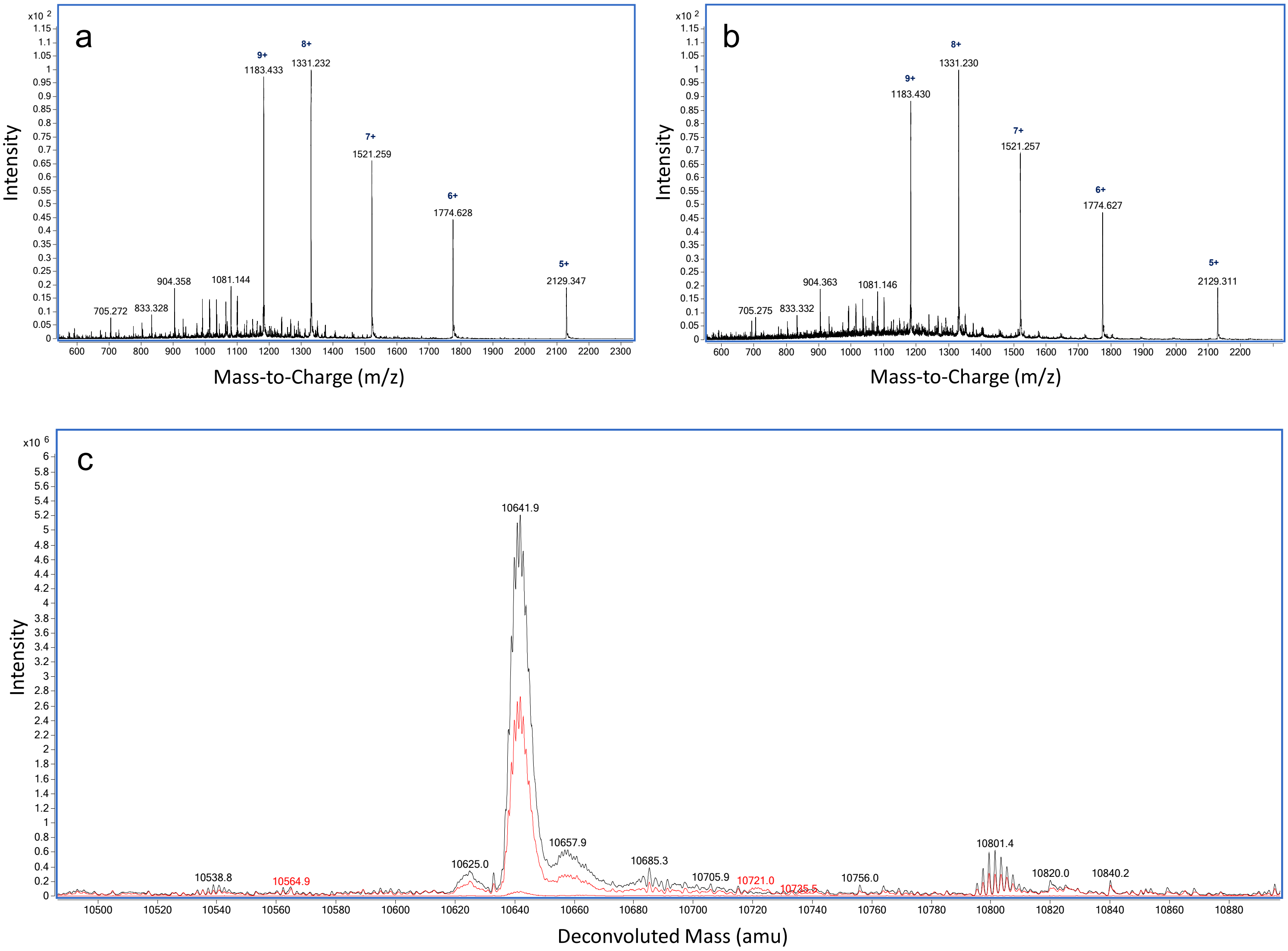


**Figure S7:** LC-MS analysis of ReoM T7A.
ESI-QTOF raw data for A) unmodified ReoM T7A and B) ReoM T7A following incubation with PrkA-KD. C) Overlaid deconvoluted mass spectrum demonstrating the lack of phosphorylation of ReoM T7A in the absence (black) and presence of PrkA-KD (red).

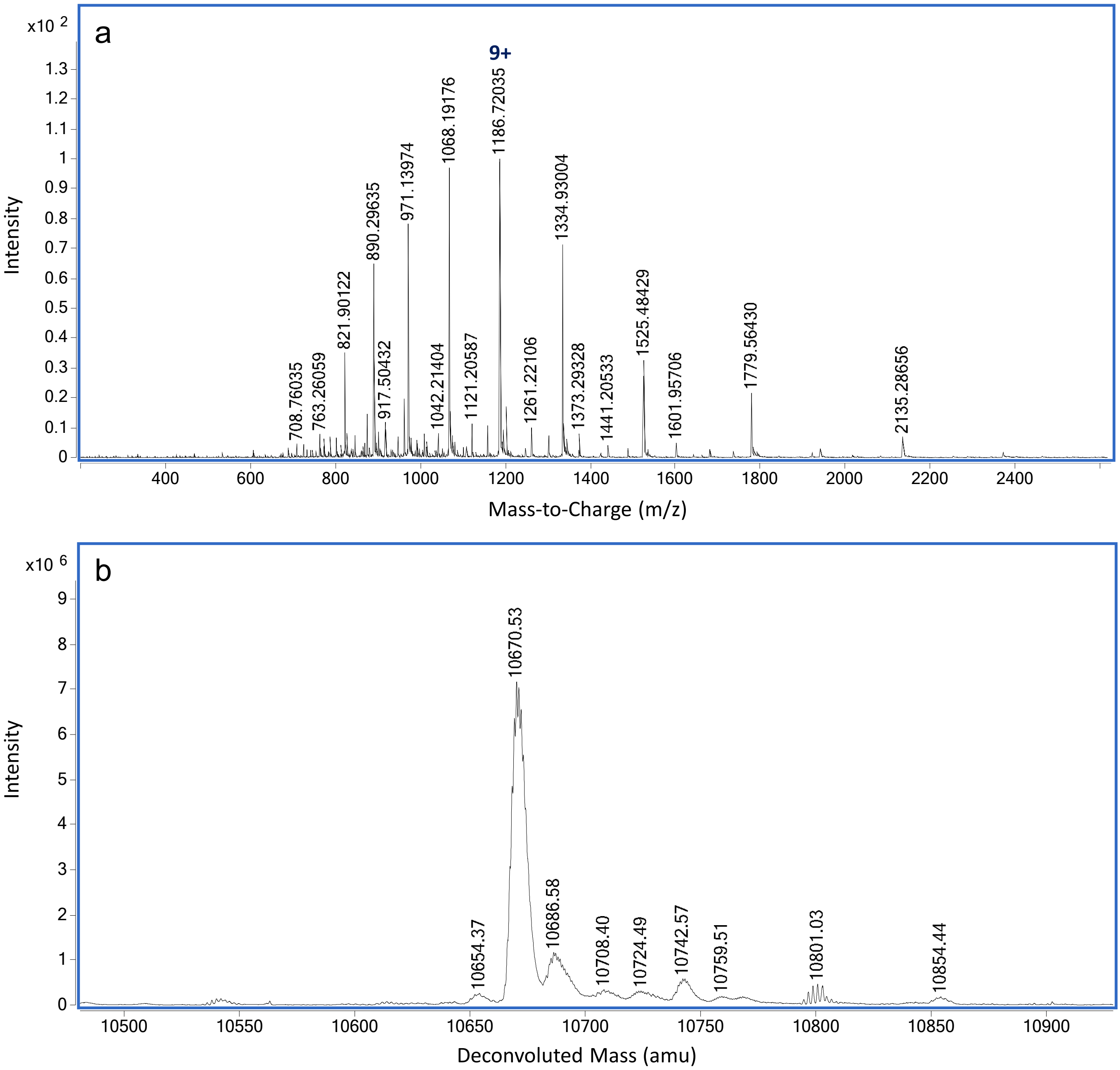


**Figure S8**: Dephosphorylation of P-ReoM by PrkP.
LC-MS analysis of ReoM after PrkP incubation in the presence of manganese. (A) ESI-QTOF raw data and (B) deconvoluted mass spectrum demonstrating the presence of non-phosphorylated ReoM.

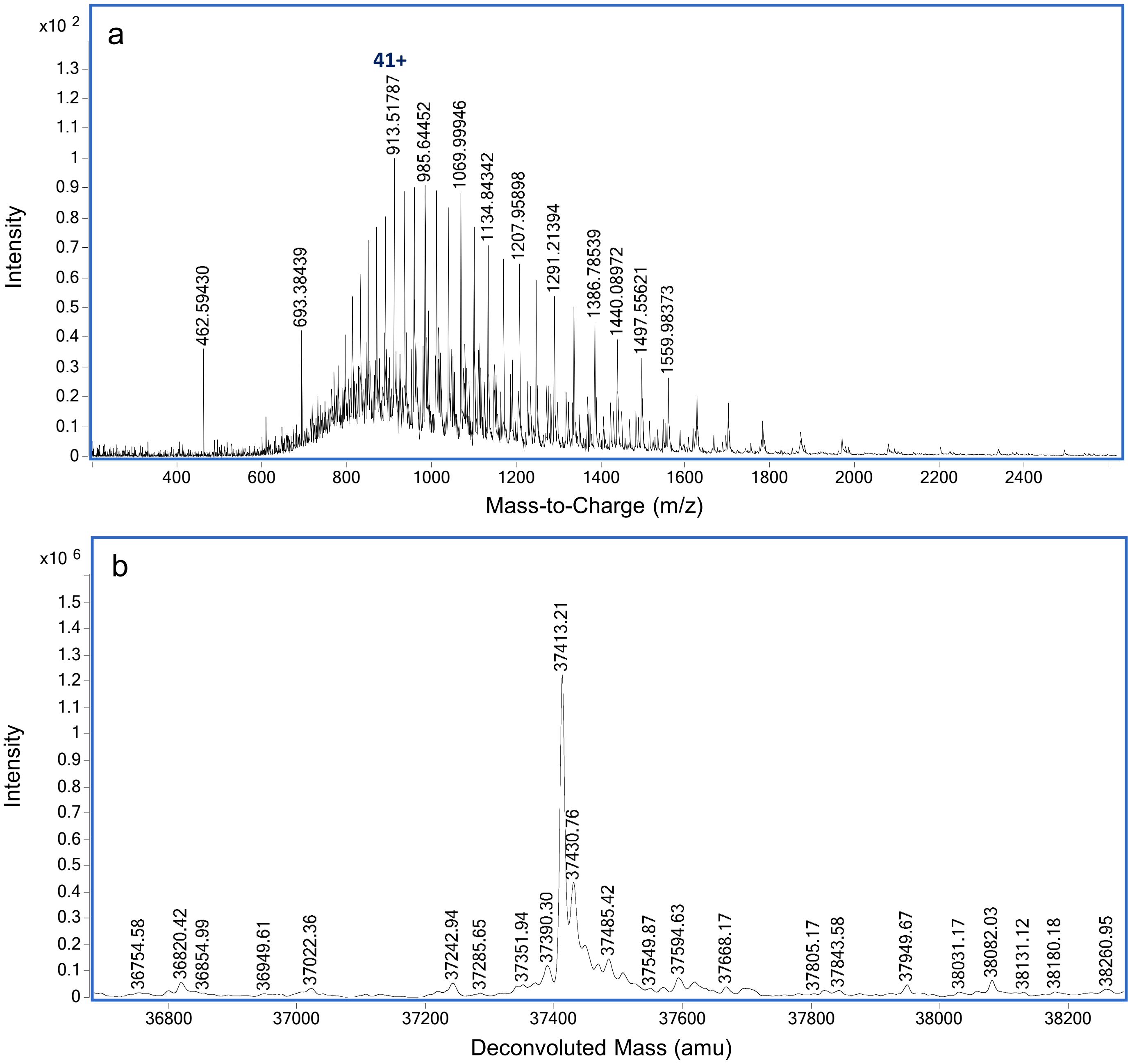


**Figure S9**: Dephosphorylation of P-PrkA-KD by PrkP.
LC-MS analysis of intact PrkA-KD following incubation with PrkP (A) ESI-QTOF raw data and (B) deconvoluted mass spectrum indicating the presence of non-phosphorylated PrkA-KD.

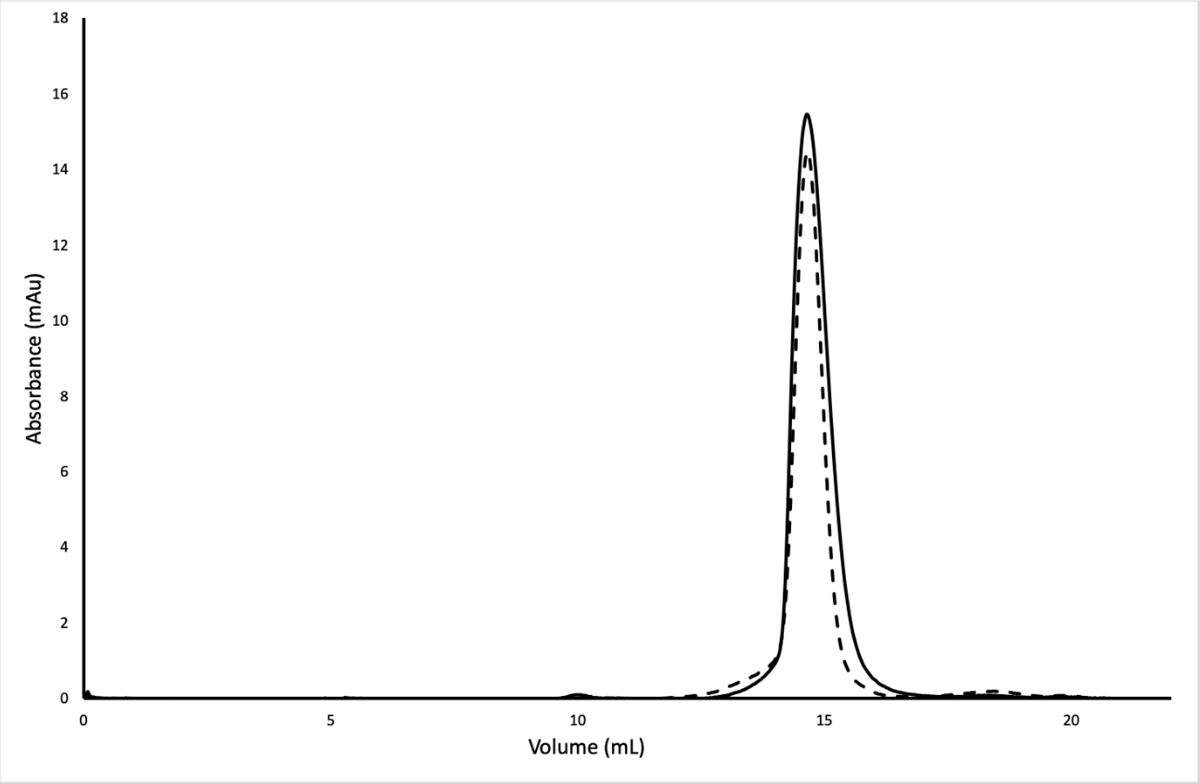


**Figure S10:** ReoM and P-ReoM have the same oligomeric state.
Size exclusion chromatography of ReoM and P-ReoM showed that phosphorylation of ReoM does not alter its oligomeric state. Both ReoM (solid) and P-ReoM (dashed) had an elution volume at 14.6 mL.

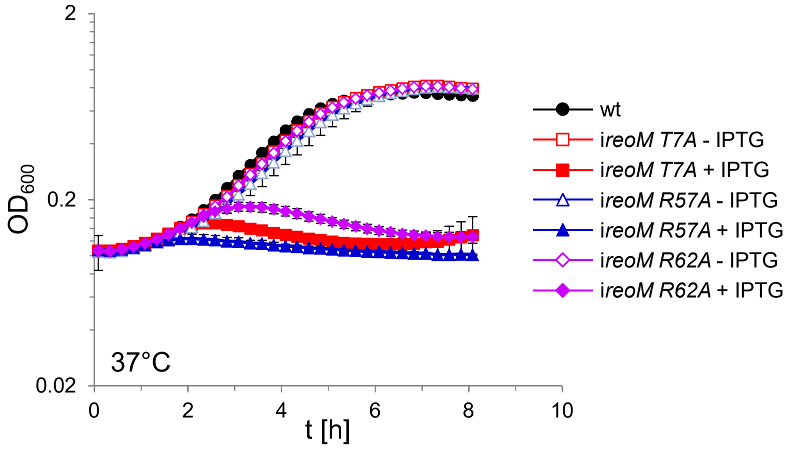


**Figure S11**: Lethality of ReoM R57A and R62A subtitutions.
*L. monocytogenes* strains EGD-e (wt), LMSW52 (i*reoM T7A*), LMSW125 (i*reoM R57A*), and LMSW126 (i*reoM R62A*) were grown in BHI broth ± 1 mM IPTG at 37°C. The experiment was repeated three times and average values and standard deviations are shown.

Wamp *et al.*, Fig. S11

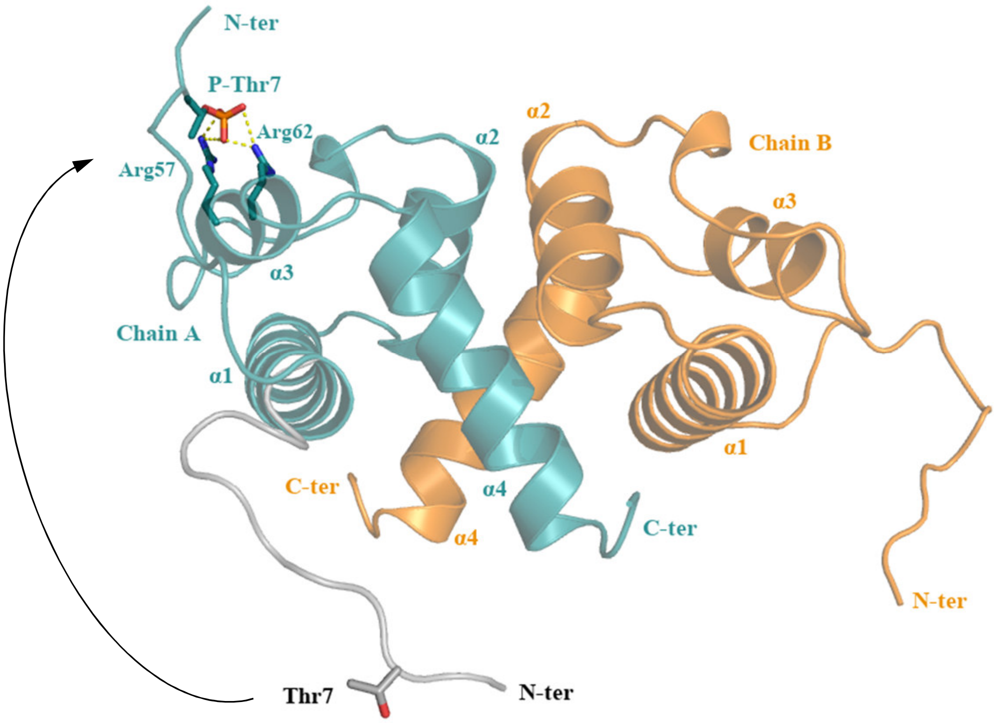


**Figure S12**: A possible conformational change of the flexible ReoM N-terminus induced by phosphorylation.
The N-terminus of ReoM might undergo a substantial movement after phosphorylation that would be stabilised in its new conformation by electrostatic interactions between the negatively charged phosphate of Thr7~P and the positively charged Arg57/Arg62 pair.

Wamp *et al.*, Fig. S12

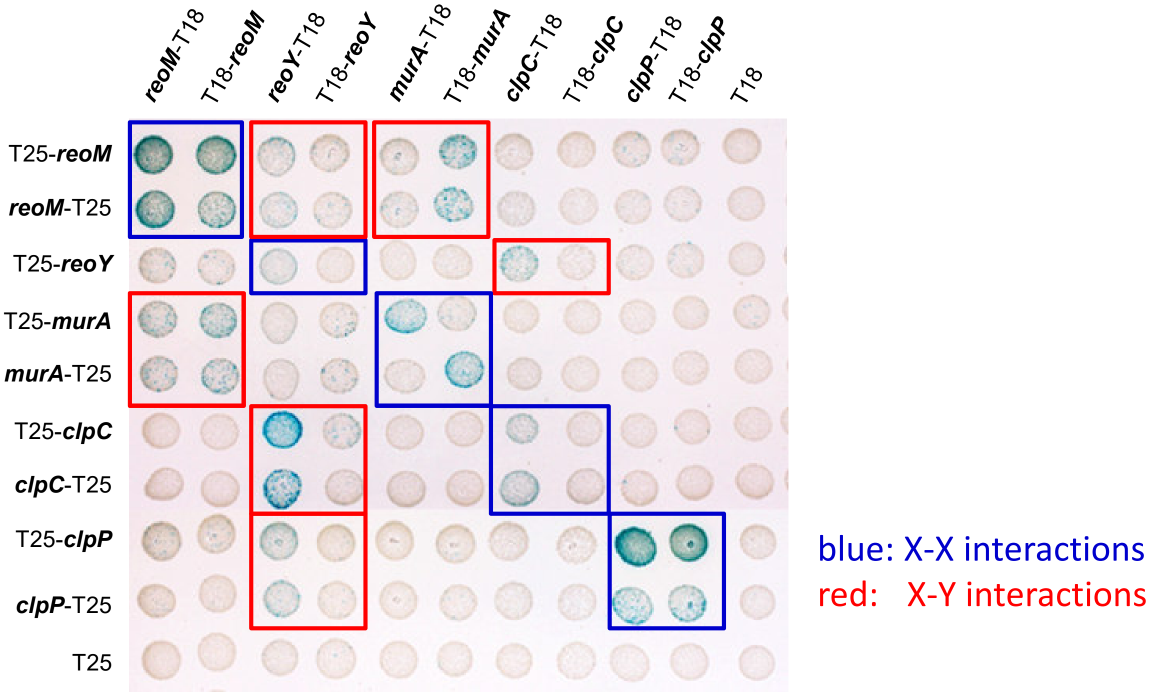


**Figure S13**: Bacterial two hybrid experiment showing interactions between MurA, ReoM, ReoY, ClpC and ClpP.
Plasmids carrying fusions of the T18- and T25-fragments of *Bordetella pertussis* adenylate cyclase fused to MurA, ReoM, ReoY, ClpC and ClpP of *L. monocytogenes* were cotransfomed into *E. coli* BTH101 and plated on selective LB agar plates containing X-Gal. Formation of blue colonies indicates interaction between the tested proteins. An illustration summarizing all detected protein-protein interactions is shown in Fig. 7C. Please note that all *murZ* fusions and the *reoY*-T25 fusion were not functional and were therefore not included.
